# A cell competition-based drug screen identifies a novel compound that induces dual c-Myc depletion and p53 activation

**DOI:** 10.1101/2020.05.16.089755

**Authors:** Dagim Shiferaw Tadele, Joseph Robertson, Richard Crispin, Maria C. Herrera, Marketa Chlubnova, Laure Piechaczyk, Pilar Ayuda-Durán, Sachin Kumar Singh, Tobias Gedde-Dahl, Yngvar Fløisand, Jørn Skavland, Jørgen Wesche, Bjørn-Tore Gjertsen, Jorrit M. Enserink

## Abstract

BCR-Abl is a driver oncogene that causes chronic myeloid leukemia and a subset of acute lymphoid leukemias. Although tyrosine kinase inhibitors provide an effective treatment for these diseases, they generally do not kill leukemic stem cells. Leukemic stem cells are cancer-initiating cells that compete with normal hematopoietic stem cells for the bone marrow niche. Using BCR-Abl as a model oncogene, we performed a drug screen based on competition between isogenic untransformed cells and BCR-Abl-transformed cells, and identified several compounds that selectively target BCR-Abl-transformed cells. Systems-level analysis of one of these novel compounds, DJ34, revealed that it induced depletion of c-Myc and activation of p53. c-Myc depletion occurred in a wide range of tumor types, including leukemia, lymphoma, lung, glioblastoma and breast cancer. Further analyses revealed that DJ34 interferes with c-Myc synthesis at the level of transcription, and we provide data showing that DJ34 is a DNA intercalator and topoisomerase II inhibitor. Physiologically, DJ34 induced apoptosis, cell cycle arrest and cell differentiation, and primary leukemic stem cells were particularly sensitive to DJ34. Taken together, we have identified a novel compound that dually targets c-Myc and p53 in a wide variety of cancers, and with particularly strong activity against leukemic stem cells.

## Introduction

BCR-Abl is the driver mutation for chronic myeloid leukemia (CML) and is also found in 25-30% of adult acute lymphoid leukemia (Ph+ ALL) ^1,2^. CML and Ph+ ALL are effectively treated with tyrosine kinase inhibitors (TKIs), such as imatinib. However, development of TKI resistance remains an issue, survival of Ph+ ALL patients is suboptimal, and new treatment strategies are required ^3^.

BCR-Abl activates downstream signaling pathways that promote cell survival and proliferation and that inhibit differentiation, such as the Ras-MAPK and PI3K-Akt pathways ^4^. Another critical component of the BCR-Abl network is the transcription factor c-Myc ^5,6^, which regulates genes important for proliferation and survival and which is important for many types of hematological and solid cancers ^7^.

Populations of self-renewing leukemic stem cells (LSCs) generate the bulk of leukemic cells ^8^. LSCs are relatively resistant to chemotherapy and persist as a potential source of relapse, and drugs that eradicate LSCs may provide durable remission. LSCs do not require BCR-Abl activity and are therefore resistant to imatinib ^9^. However, they are highly reliant upon both c-Myc and p53, and dual targeting of both c-Myc and p53 has been shown to selectively and synergistically eliminate LSCs ^6^, demonstrating a need for novel compounds that simultaneously inhibit c-Myc and activate p53.

LSCs compete with healthy hematopoietic stem cells (HSCs) for the bone marrow niche, which constitutes a functional vulnerability of primitive leukemia cells ^10^. However, cell competition is not typically assayed in high-throughput drug screens, and drugs that reduce the competitiveness of cancer cells without directly affecting cell viability are likely to be discarded.

We reasoned that a straightforward cell-competition assay, in which healthy cells and isogenic oncogene-expressing cells compete against each other, would efficiently identify novel compounds that selectively target oncogene-transformed cells. We identified several compounds that preferentially inhibit BCR-Abl-transformed cells, including a compound with anti-LSC activity that dually targets c-Myc/p53.

## Results

### High-throughput cell competition drug screen for BCR-Abl-expressing cells

To identify novel compounds that target BCR-Abl-driven leukemias, we developed an isogenic cell competition-based drug screen by stably transfecting Ba/F3 cells with BCR-Abl. Transfection of Ba/F3 cells with BCR-Abl transformed the cells and resulted in IL3-independent growth (Fig. 1A). Consistent with previous reports ^17,18^, BCR-Abl-transformed cells were significantly more sensitive to imatinib than wild-type (WT) cells, demonstrating they had become oncogene-addicted (Fig. 1A). Next, BCR-Abl-expressing cells were stably transfected with EGFP, and WT cells with mCherry. These cells were mixed and treated with imatinib for 72 h, after which the BCR-Abl:WT cell ratio was measured by flow cytometry (Fig. 1B). Imatinib conferred a competitive disadvantage upon BCR-Abl cells (Fig. 1C). Importantly, the competition assay was significantly more sensitive than the commonly used CellTiter-Glo cell viability assay (Fig. 1D).

**Figure 1.**
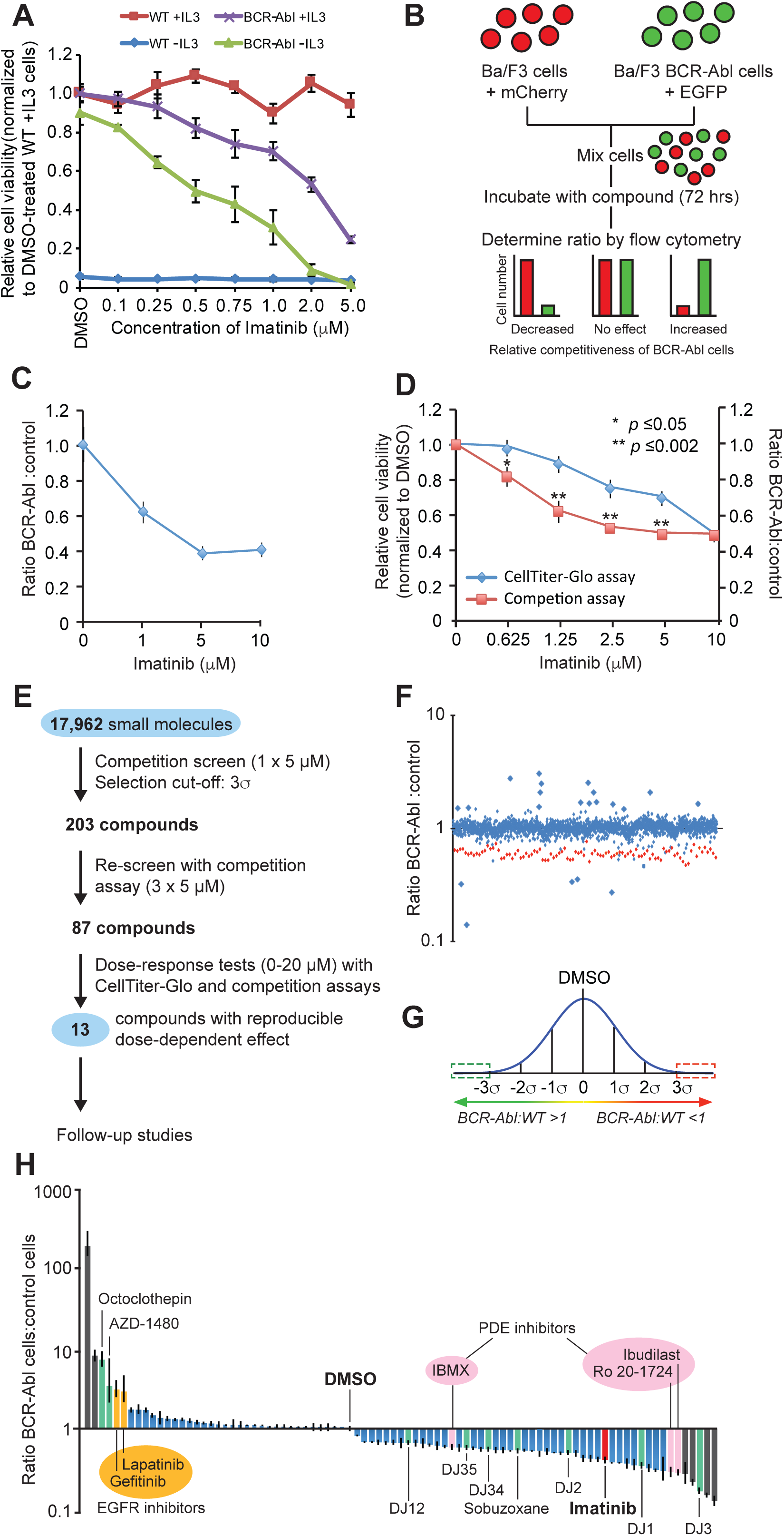
Development of an isogenic, cell competition-based drug screen to identify compounds that modulate the competitiveness of BCR-Abl-expressing cells. (**A**) Effect of imatinib on the relative viability of wild-type (WT) or BCR-Abl-expressing Ba/F3 cells. (**B**) Schematic overview of the competition-based drug assay. (**C**) Ratio of BCR-Abl-expressing cells compared to WT cells following treatment with imatinib for 72 hrs. (**D**) Comparison of the selective targeting of BCR-Abl cells by imatinib as measured using a cell viability assay or the competition assay. (**E**) Schematic overview of the different stages of the competition-based drug screen. (**F**) Effect of different drugs on the competitiveness of BCR-Abl cells. Red dots represent imatinib-treated cells. For display purposes, only a selection of tested compounds is displayed (approx. 20%). (**G**) Compounds that fell outside the 3σ interval (compared to DMSO-treated cells) were selected for follow-up. (**H**) The 87 compounds that were considered for follow-up studies after re-screening of each compound in triplicate, but prior to dose-response tests (see panel E). All values were normalized to DMSO. Grey bars represent compounds that were non-replicable and therefore removed. Green bars represent compounds with a reliable dose-dependent effect, which were selected for follow-up studies; blue bars represent compounds with reliable dose-dependent effects but that were not selected for follow-up experiments, mainly due to issues with availability and pricing. Error bars in the relevant panels indicate standard deviation and statistical significance was determined using Student’s t-test.

Using the competition assay we screened 17,962 compounds for selective inhibition of BCR-Abl-expressing cells (Fig. 1E). Most compounds had little or no impact on the BCR-Abl:WT cell ratio (Fig. 1F). The effect of the drugs followed a Gaussian distribution, and 203 compounds were selected that fell outside the 3σ interval (Fig. 1G). These compounds were re-tested three times, resulting in 87 drugs that significantly (*p*<0.05) altered the BCR-Abl:WT cell ratio (Fig. 1H). This included imatinib, thus validating the screen. All 87 compounds were subjected to dose-response tests with freshly prepared drug stocks using competition assays and viability assays. We found that some compounds did not replicate the effect of the original library compounds (grey bars in Fig. 1H). This may have been due to errors during preparation of the original library, contaminants in the initial screen, or compound degradation products. Non-reproducible compounds and compounds that were excessively expensive to synthesize were discarded, leaving a total of 13 compounds that either increased (four compounds, although high doses induced general toxicity, see Suppl. Fig. S1A) or decreased (nine compounds; Fig. 2) the competitiveness of BCR-Abl-expressing cells.

**Figure 2.**
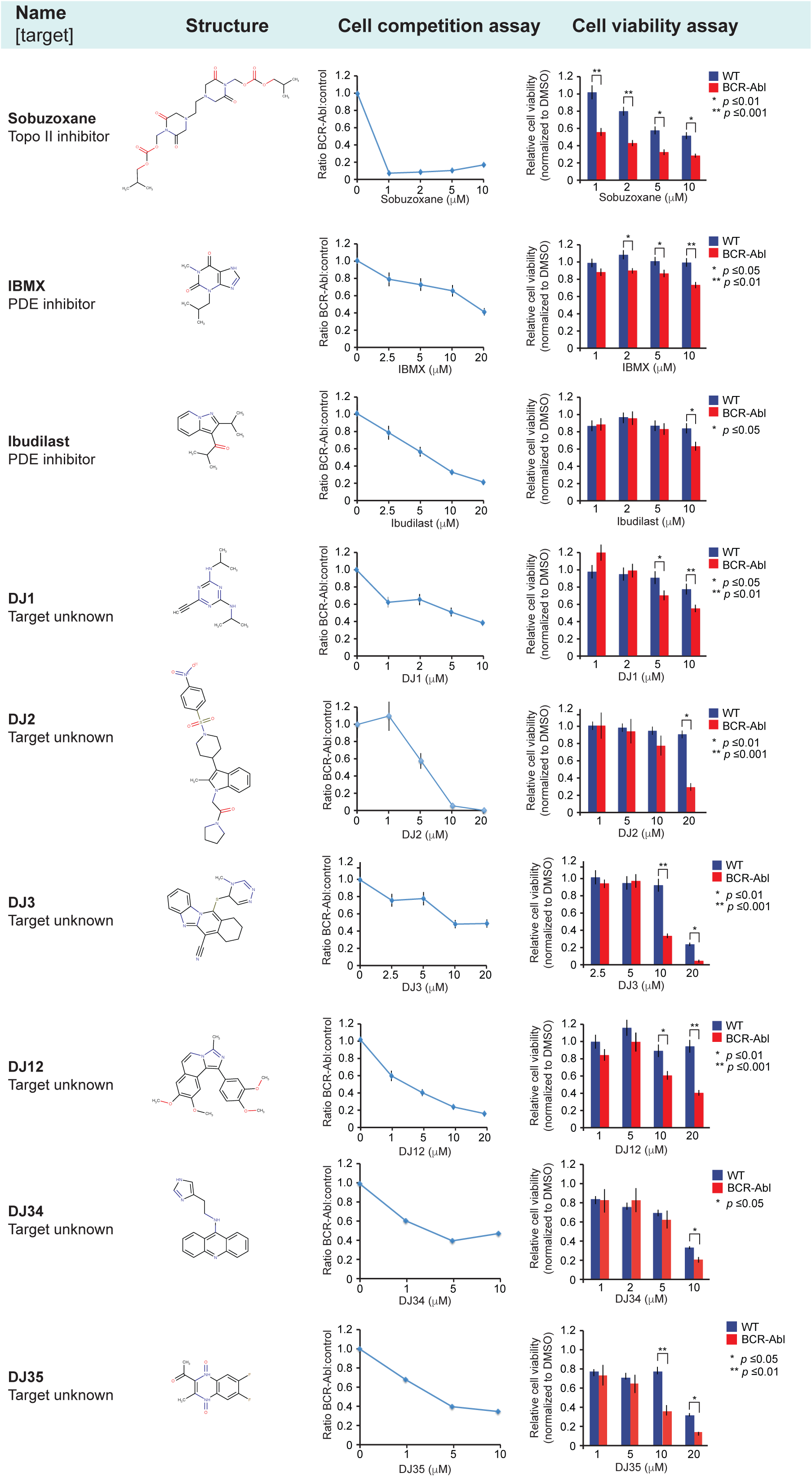
Nine compounds that decreased the relative competitiveness of BCR-Abl-expressing cells compared to WT cells. Overview of the names, known molecular targets and structures of the compounds, as well as the results obtained with the cell competition and cell viability assays. All values were normalized to DMSO. Error bars indicate standard deviation and statistical significance was determined using Student’s t-test.

### Compounds that increase the competitiveness of BCR-Abl-expressing cells

Among the four compounds that promoted relative competitiveness of BCR-Abl cells was the JAK2 inhibitor AZD1480 (Suppl. Fig. S1A). Structurally unrelated JAK2 inhibitors had the same effect (Suppl. Fig. S1B), while co-treatment with imatinib restored the dependency of BCR-Abl-expressing cells to JAK2 (Suppl. Fig. S1C), consistent with previous studies ^19^.

The EGFR inhibitors lapatinib and gefitinib and the G protein-coupled receptor (GPCR) inhibitor octoclothepin also promoted the competitiveness of BCR-Abl cells (Suppl. Fig. S1A), although the reasons for this are currently unclear. Ba/F3 cells do not normally express EGFR, suggesting an off-target effect of EGFR inhibitors, e.g. by inhibiting JAK kinases ^20^. Octoclothepine inhibits GPCRs that elevate intracellular cAMP levels ^21^, and cAMP impairs survival of BCR-Abl-expressing cells (see below).

### Compounds that reduce the competitiveness of BCR-Abl-expressing cells

We identified nine compounds that reduced the competitiveness of BCR-Abl cells, including Sobuzoxane (topoisomerase II inhibitor) and the PDE inhibitors IBMX and Ibudilast (Fig. 2). Their effect was validated by unrelated inhibitors (Suppl. Fig. S2A-E). PDE inhibitors increase cellular cAMP levels, which decreases the growth rate of multiple tumor cell types by activating PKA ^22^. The cAMP synthesis-activating agent forskolin ^22^ also inhibited BCR-Abl cells, whereas the inactive forskolin analog dideoxyforskolin had no effect (Suppl. Fig. S2F,G). Similar results were obtained with the PKA agonist 8Br-cAMP, but not with a cAMP analog that cannot activate PKA ^16^ (Suppl. Fig. S2H,I). These data demonstrate that drugs that increase cAMP levels selectively inhibit BCR-Abl-expressing cells.

Six uncharacterized compounds selectively inhibited BCR-Abl-expressing cells (DJ1, DJ2, DJ3, DJ12, DJ34 and DJ35; Fig. 2), but they did not directly target BCR-Abl (Suppl. Fig. S3). Several of these compounds also inhibited the human CML cell lines MEG-01, KU-812 and K562, as well as the human Ph+ ALL cell line SD-1, which was not effectively killed by imatinib (Suppl. Fig. S4), indicating that these compounds may serve as a starting point for development of alternative forms of therapy for both CML and Ph+ ALL patients.

### DJ34 selectively kills BCR-Abl-positive leukemia cells

We analyzed the drug-like properties of the novel compounds using SwissADME ^23^ and found that DJ2, DJ12 and DJ35 have unfavorable drug-like properties, whereas DJ1 and DJ3 contain an alkyne group and a nitrile group, respectively, which can be unstable and chemically reactive *in vivo*. DJ34 was predicted to have excellent drug-like properties (Suppl. Table S2, tab 1), which was confirmed by initial ADME-PK analyses (Suppl. Table S2, tab 2-6). We confirmed that freshly synthesized DJ34 targeted BCR-Abl-transformed Ba/F3 cells more efficiently than the isogenic parental cells (Fig. 3A). DJ34 also killed primary cancer cells derived from a Ph+ ALL patient more efficiently and at lower doses than imatinib (Fig. 3B). Importantly, blast cells derived from ALL and mixed B-ALL/AML patients were more sensitive to DJ34 than healthy bone marrow cells (Fig. 3C), suggesting a therapeutic window.

**Figure 3.**
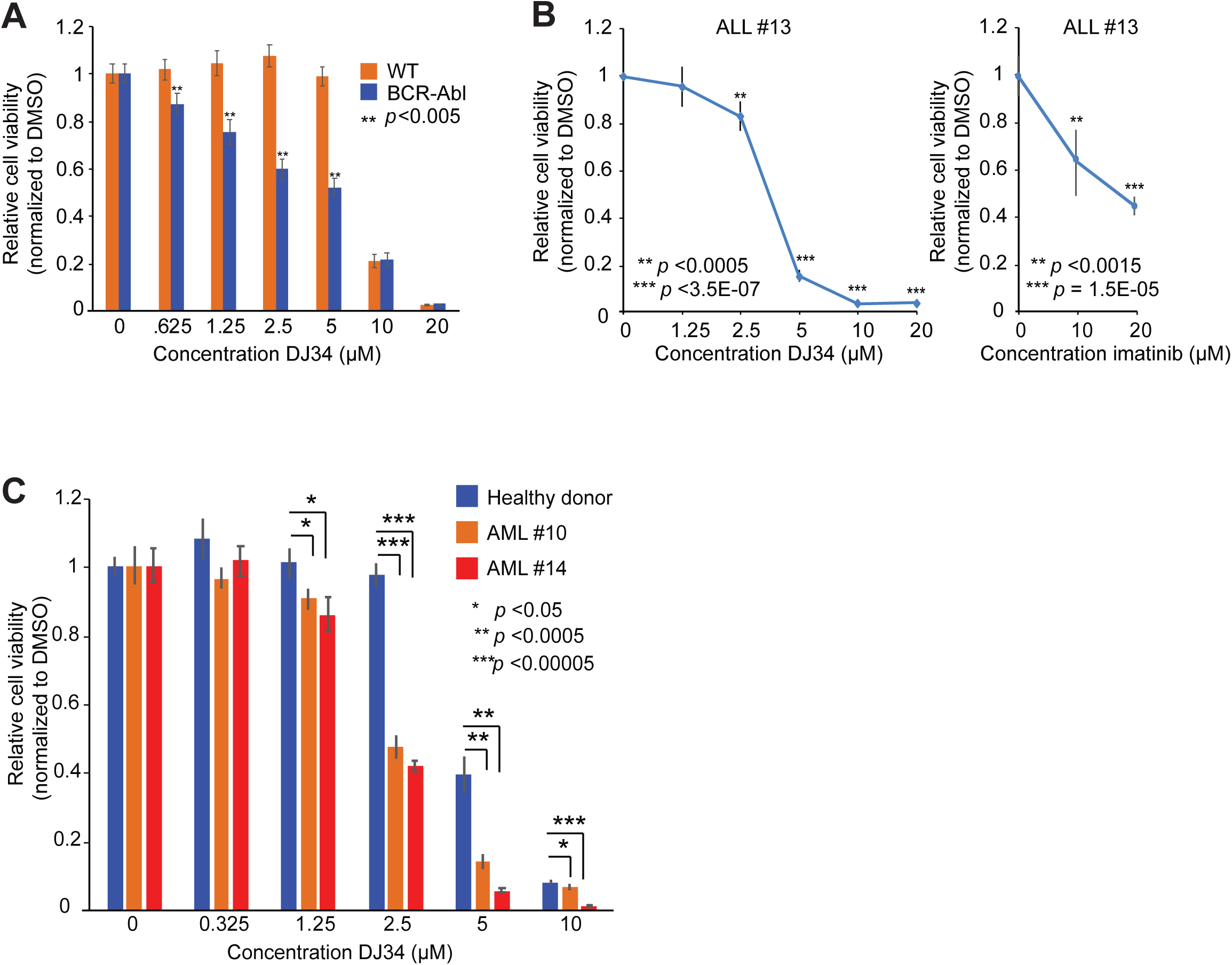
DJ34 selectively targets BCR-Abl+ leukemia cells. **(A)** BCR-Abl-transformed Ba/F3 cells are significantly more sensitive to DJ34 than isogenic control cells. Cells were incubated with increasing concentrations of DJ34 for 72 hrs, after which cell viability was analyzed by CellTiterGlo. **(B)** DJ34 efficiently kills primary Ph+ ALL cells. ALL cells were isolated from the bone marrow from a Ph+ ALL patient and incubated for 72 hrs with the indicated concentrations of DJ34 (*left*) or imatinib (*right*), after which cell viability was analyzed as in panel *A*. (**C**) Primary ALL cells are more sensitive to DJ34 than bone marrow cells derived from a healthy donor. Cells were isolated from bone marrow samples of an ALL patient (patient #10), a mixed B-ALL/AML patient (patient #14) and a healthy donor and incubated for 72 hrs with the indicated concentrations of DJ34, after which relative cell viability was analyzed by CellTiterGlo. Error bars indicate standard deviation and statistical significance was determined using Student’s t-test.

### DJ34 inhibits the c-Myc transcriptional program and activates the p53 program

To better understand how DJ34 may inhibit cancer cells, we first performed phosphoflow cytometry experiments to investigate its effect on a panel of oncogenic signaling pathways. DJ34 had no effect on any of the examined phosphoproteins (Suppl. Fig. S5A). AKT pS473 and STAT5 pY694, which depend on BCR-Abl signaling, were also not affected by DJ34 (Suppl. Fig. S5B), consistent with our observations that DJ34 does not directly inhibit BCR-Abl (Suppl. Fig. S3). This shows that DJ34 does not affect several canonical oncogenic pathways.

We then decided to apply a broader, unbiased approach to determine how DJ34 may affect cells using a combination of RNA sequencing (RNA-seq) and mass spectrometry (MS)-based phosphoproteomics for multi-parameter analysis of the cellular programs altered by DJ34 (Fig. 4A). RNA-seq analysis identified 1206 and 1705 gene transcripts that were at least two-fold decreased or increased in abundance by DJ34 treatment, respectively (Suppl. Table S3). Gene set enrichment analysis (GSEA) revealed that c-Myc-dependent genes were significantly enriched in the RNA-seq dataset, as well as genes associated with the p53 pathway (Fig. 4B). Closer inspection confirmed that DJ34 down-regulated genes such as *EIF4, CD47, CDC2, CCND2* and *RCC1*, which are activated by c-Myc (e.g. ^24^; Fig. 4C, *upper panel*). Conversely, treatment with DJ34 promoted the expression of genes that are inhibited by c-Myc, such as *NDRG1, CDKN1A* and *GADD45* (^24,25^; Fig. 4C, *lower panel*). Similar findings were obtained by RT-qPCR analysis of several c-Myc target genes (Fig. 4D). Gene Ontology (GO) analysis of the genes up-regulated by DJ34 revealed a significant enrichment of terms associated with reduced c-Myc function and/or increased p53 signaling, such as ‘mitotic cell cycle’, ‘positive regulation of programmed cell death’ and ‘positive regulation of cell differentiation’ (Suppl. Fig. S6A).

**Figure 4.**
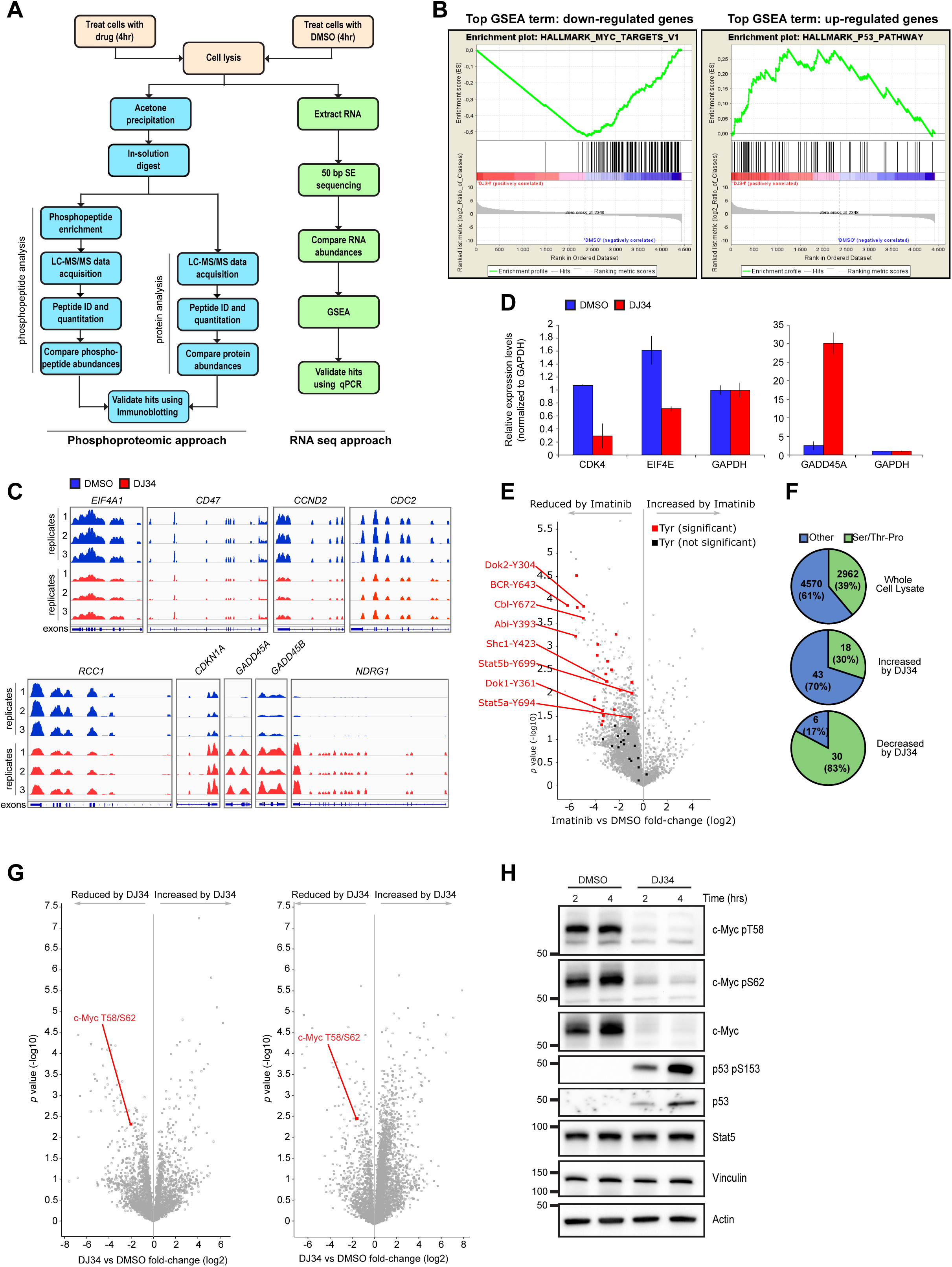
Multi-parameter analysis to determine the effect of DJ34 on transcription and cell signaling. (**A**) Workflow for the combined phosphoproteomic and RNA sequencing analysis of BCR-Abl-expressing cells treated with DJ34. (**B**) GSEA of significantly up- and down-regulated transcripts following RNA-seq, showing the top scoring enriched gene set terms. (**C**) Integrative genomics view (IGV) of the mRNA levels of selected c-Myc target genes. Y axes show RFPKM values. Range of the respective y axes: *EIF4A1*: 0-1500; *CD47*: 0-1500; *CCND2*: 0-2000; *CDC2*: 0-1000; *RCC1*: 0-1000; *CDKN1A*: 0-4500; *GADD45A*: 0-200; *GADD45B*: 0-400; *NDRG1*: 0-1000. (**D**) RT-qPCR validation of selected c-Myc target genes after treatment with DJ34. Error bars, standard deviation. (**E**) Volcano plot showing phosphosites modulated by imatinib. (**F**) Proportions of proline-directed phosphophorylation sites affected by DJ34. (**G**) Volcano plots of two biological repeats showing phosphosites regulated by DJ34 treatment. (**H**) Immunoblot analysis of BCR-Abl-expressing Ba/F3 cells treated with DMSO or DJ34 for 2 and 4 hrs.

Phosphoproteomic analysis of cells treated with imatinib or DJ34 indicated that these compounds have distinct effects on cellular signaling pathways. We identified a similar number of unique phosphopeptides from DMSO-, imatinib- and DJ34-treated cells (Suppl. Fig. S6B). Whilst imatinib decreased the abundance of several tyrosine-phosphorylated peptides, including sites previously associated with BCR-Abl signaling (thus validating our approach; Fig. 4E; Suppl. Table S4), tyrosine phosphosites did not appear to be affected by DJ34 treatment (Suppl. Table S5). Instead, a high proportion of the phosphosites reduced by DJ34 were proline-directed phosphosites (Fig. 4F), including the residues Thr58 and Ser62 on c-Myc (Fig. 4G), consistent with our RNA-seq data indicating that DJ34 inhibits c-Myc activity (Fig. 4B-D).

Validation of RNA-seq and MS data by immunoblotting confirmed that phosphorylation of c-Myc Thr58 and Ser62 was decreased by DJ34, but also showed that total c-Myc protein levels were strongly reduced (Fig. 4H; total c-Myc levels were not detected in by MS, most likely because it is a low-abundance protein that could only be detected after enrichment of phosphorylated peptides). Furthermore, while p53 was undetectable in the lysates of DMSO-treated cells, DJ34 treatment increased total p53 levels as well as Ser15-phosphorylated p53 (a marker of active p53; Fig. 4H). Together, these data indicate that DJ34 simultaneously inhibits c-Myc and activates p53.

### DJ34 induces depletion of c-Myc in a wide variety of tumor types

A compound that both inhibits c-Myc while stabilizing p53 could provide a valuable therapeutic agent not only for leukemia but for a wide range of cancers ^6,26-28^. We found that DJ34 treatment resulted in c-Myc depletion in all cancer cell lines tested, including glioblastoma, breast cancer, and lung cancer cell lines (Fig. 5). DJ34 treatment depleted c-Myc in primary human AML blasts as well (Fig. 5), showing that the anti-c-Myc effect of DJ34 is not restricted to laboratory cell lines. Furthermore, DJ34 treatment increased p53 levels for almost all cancer cell lines that exhibited low p53 levels under control conditions (Fig. 5). This included the lung cancer cell lines A-549 and H-460, the breast cancer cell line MCF-7, and the glioblastoma cell line U-87, all of which express wild-type p53 ^29^. The same was observed for the leukemia cell line SD-1 (unknown p53 status), whereas p53 was undetectable in the p53-mutant CML cell line K562 ^30^. Two other p53-mutant cell lines, H-1975 (R273H gain of function mutation) and HCC-827 (V218DEL) ^31,32^, exhibited high basal p53 levels that were not increased by DJ34. Together, these data show that DJ34 treatment broadly inhibits c-Myc levels while at the same time activating p53.

**Figure 5.**
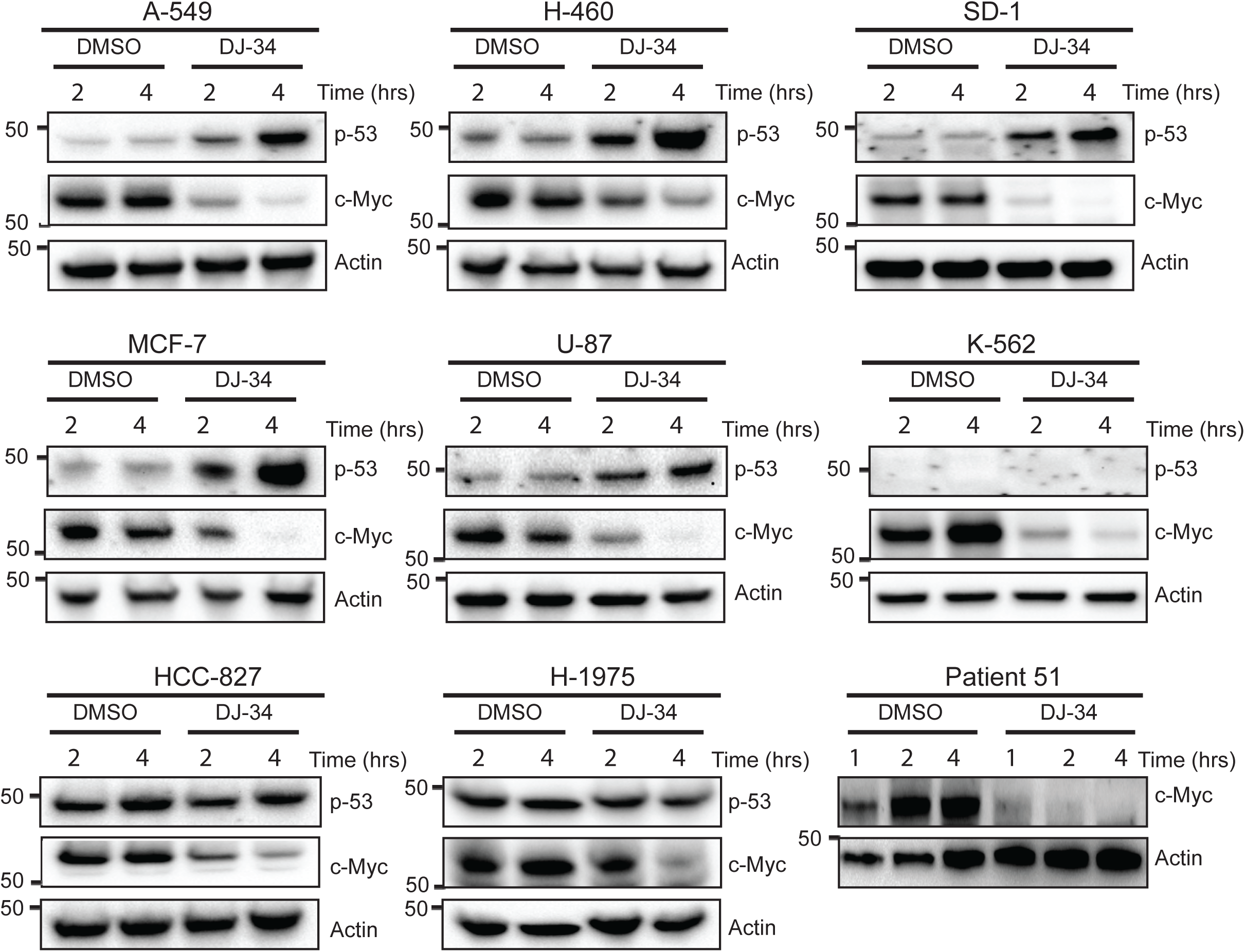
DJ34 targets c-Myc and p53 in a variety of cancer cells. Immunoblotting analysis for c-Myc, p53 and Actin in various cancer cell lines treated with DMSO or 10 µM DJ34 for 2 and 4 hrs. In addition, bone marrow cells derived from an AML patient were treated with DMSO or DJ34 for 1, 2 and 4 hrs and c-Myc and Actin levels were analyzed by immunoblotting (“Patient 51”, bottom right panel).

### DJ34-induced cellular depletion of c-Myc requires an intact STUB1/CHIP pathway

We wished to better understand the cellular requirements for the DJ34-induced reduction in c-Myc. The proteasomal inhibitor MG132 prevented DJ34-induced c-Myc depletion (Suppl. Fig. S7A), suggesting that DJ34 treatment leads to proteasomal depletion of c-Myc. In many cancers, proteasomal degradation of c-Myc is prevented by increased phosphorylation of Ser62. This can occur via either hyperactive proline-directed kinases including MAPKs and CDKs or inactivation of the phosphatase PP2A ^33^. DJ34 primarily down-regulated proline-directed phosphorylation sites (Fig. 4F), so we assessed whether DJ34 induces proteasomal depletion of c-Myc via regulating its phosphorylation status. *In vitro* kinase analysis using the KINOME*scan* platform (www.discoverx.com) revealed that DJ34 had moderate or no effect on the activity of 55 kinases (Suppl. Table S6), including the *bona fide* c-Myc kinases ERK and Cdk2. In addition, pretreatment with the potent PP2A inhibitor okadaic acid did not prevent DJ34-induced depletion of c-Myc (Suppl. Fig. S7B; phosphorylation of the direct PP2A target Akt is shown as control ^34^). These data indicate that DJ34 does not cause c-Myc depletion by targeting kinases or phosphatases known to regulate c-Myc proteasomal degradation.

We next tested the effect of DJ34 on other proteins known to regulate c-Myc proteasomal degradation. Certain Skp, Cullin, F-box (SCF) complexes can ubiquitinate c-Myc to target it for destruction ^33^. SCF activity requires prior neddylation of the Cullin RING ligase subunit by the ubiquitin-like protein Nedd8 ^35^. SCF activity can be efficiently inhibited by the small molecule inhibitor MLN4924, which targets the E1 Nedd8 activating enzyme ^36^. While pretreatment of cells with MLN4924 considerably increased c-Myc levels, DJ34 treatment still resulted in depletion of a large proportion of c-Myc (Suppl. Fig. S7C). Treatment with DJ34 also induced strong c-Myc depletion in *FBXW7*^-/-^ cells (Suppl. Fig. S7D), which lack the main Cullin responsible for c-Myc degradation ^37^. We did observe that the rate of c-Myc depletion was slower in *FBXW7*^-/-^ cells than in WT cells (see graph in Suppl. Fig. S7D), suggesting that part of the c-Myc pool might be degraded by FBW7 upon DJ34 treatment. Alternatively, it is possible that in *FBXW7*^-/-^ cells the c-Myc degradation machinery is saturated due to the high levels of basal c-Myc, resulting in an apparent reduction in the rate of DJ34-induced c-Myc depletion. Consistent with the latter hypothesis, the levels of the FBW7 target cyclin E ^38^ were not affected by DJ34 (Suppl. Fig. S7E), arguing against the possibility that DJ34 directly activates the SCF^FBW7^ complex to degrade c-Myc.

Next, we focused on CHIP (also known as STUB1), which is an E3 ubiquitin ligase that mediates c-Myc depletion independently of the phosphorylation status of c-Myc. Strikingly, even after 6 hrs of DJ34 treatment, CHIP^-/-^ MEFs still had not succeeded in degrading c-Myc (Suppl. Fig. S7F), showing that an intact CHIP pathway is required for depletion of c-Myc by DJ34.

### DJ34 induces depletion of c-MYC mRNA

To further delineate the process of c-Myc depletion, we combined DJ34 with the ribosome inhibitor cycloheximide and monitored c-Myc protein levels over time. Interestingly, DJ34 did not appear to accelerate cycloheximide-induced depletion of c-Myc (Suppl. Fig. S7G), suggesting that DJ34 primarily acts upstream of the ribosome. We used qPCR to determine whether DJ34 has a pre-translational effect on c-Myc, which revealed a significant reduction in *c-MYC* mRNA levels upon DJ34 treatment in all cell lines tested, including K562, MV4-11, HCT-116 and MIA-PaCa cells (Fig. 6A). These data suggest that DJ34 inhibits transcription of *c-MYC*.

**Figure 6.**
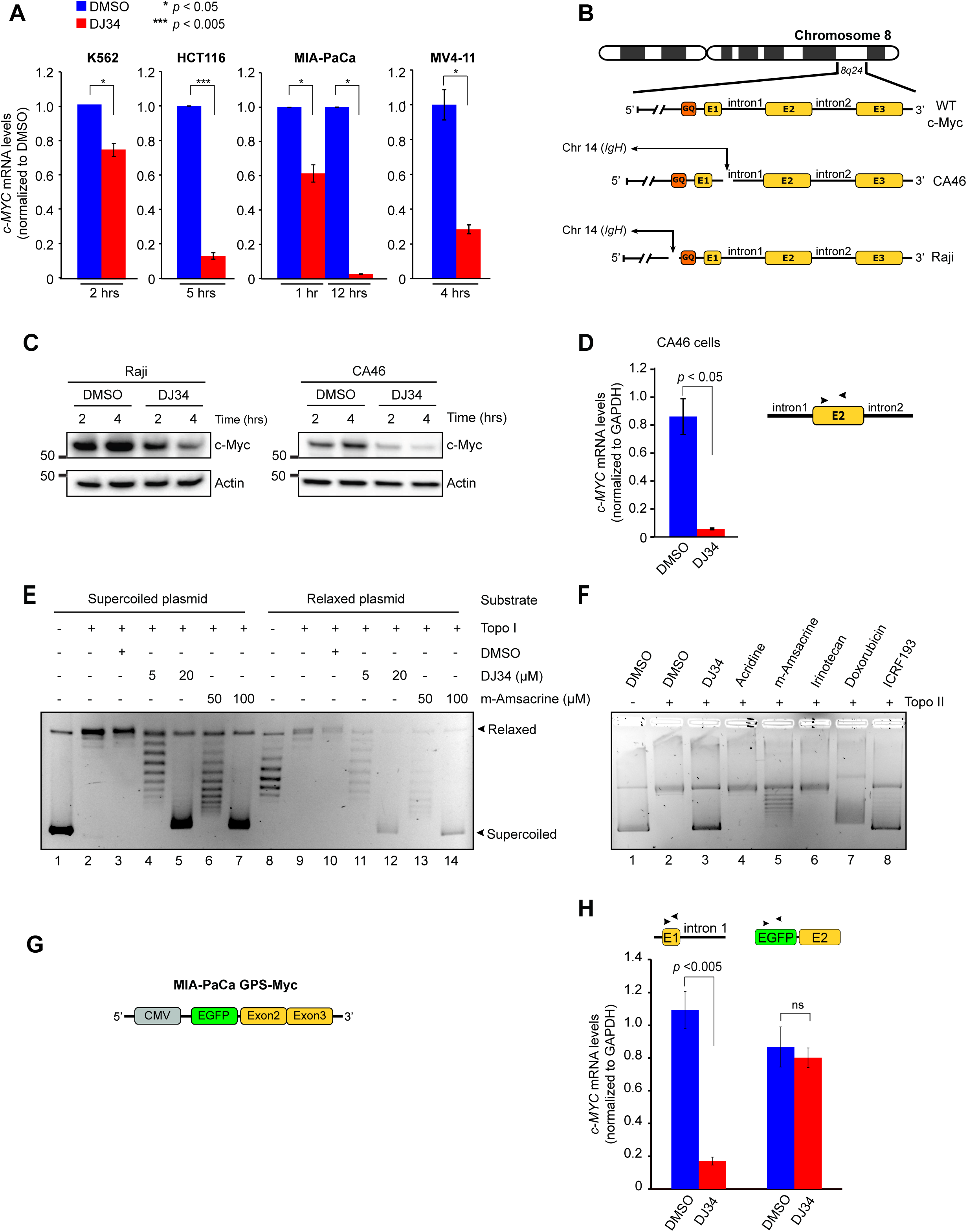
DJ34 intercalates the DNA and inhibits topo II but not topo I. (**A**) RT-qPCR analysis of *c-MYC* mRNA levels after K562, MV4-11, HCT116 and MIA-PaCa cells were treated with DMSO or 10 µM DJ34 for the indicated time points. mRNA levels were first normalized to GAPDH and then to the DMSO treatment. (**B**) Schematic overview of *c-MYC* status in non-translocated (WT) cells compared to the lymphoma cell lines CA46 and Raji, both of which possess a reciprocal translocation between *IgH* and *c-MYC*. The translocation in CA46 cells leads to loss of the G4 quadruplex, whereas in Raji cells the G4 quadruplex is retained. (**C**) Immunoblot analysis of Raji and CA46 cells treated with DMSO or 10 µM DJ34 for 2 and 4 hrs. (**D**) RT-qPCR analysis of *c-MYC* mRNA levels in CA46 cells treated with DMSO or 10 µM DJ34 for 4 hrs. Primer pairs were designed to target exon 2 (as depicted above the bar graph), which almost exclusively measures levels of the translocated *c-MYC* allele, because expression of the non-translocated allele is approximately 1000-fold lower than the translocated allele ^41^. (**E**) DNA intercalation/topo I inhibition assay showing that DJ34 intercalates the DNA but does not inhibit topo I (see Suppl. Fig. S7I for a detailed explanation of the assay). Supercoiled and relaxed plasmid substrates were incubated with different compounds and purified topo I as indicated, after which the products were analyzed by agarose gel electrophoresis. (**F**) DJ34 inhibits topo II. Supercoiled plasmid substrate was incubated with topo II and various compounds as indicated above the figure, after which the products were analyzed by agarose gel electrophoresis. All compounds were used at 10 µM. (**G**) Illustration depicting the *c-MYC* cDNA sequence randomly integrated into the genome of MIA PaCa cells (MIA-PaCa GPS-Myc). This cDNA lacks exon 1 and all introns. (**H**) RT-qPCR analysis of *c-MYC* mRNA levels after MIA-PaCa GFP-Myc cells were treated with DMSO or 10 µM DJ34 for 4 hrs. Primers were designed targeting different regions of *c-MYC* (depicted above each set of bars) to distinguish between mRNA generated from wild-type *c-MYC* and cDNA-encoded *c-MYC*.

The reduction in c*-MYC* mRNA levels by DJ34 could be due to altered mRNA stability rather than inhibition of transcription. To distinguish between these possibilities, we blocked transcription with the strong RNA polymerase II inhibitor α-amanitin ^39^. We hypothesized that if DJ34 destabilizes mRNA, combining DJ34 with α-amanitin should result in more rapid loss of c-Myc, whereas there should be no added effect if DJ34 impedes *c-MYC* transcription, since mRNA synthesis is already fully blocked by α-amanitin. While treatment with α-amanitin resulted in efficient removal of c-Myc, α-amanitin did not appear to potentiate DJ34-induced loss of c-Myc (Suppl. Fig. S7H), indicating that DJ34 and α-amanitin target the same process, i.e. *c-MYC* transcription.

We reasoned that DJ34 might inhibit *c-MYC* transcription by interacting with regulatory elements in the *c-MYC* gene, such as the G-Quadruplex in the promoter region (G4). This is a helical DNA structure formed by guanine tetrads, and several compounds can stabilize the quadruplex to inhibit *c-MYC* transcription ^40^. We examined c-Myc protein levels in two lymphoma cell lines in which genomic translocations of *IgH* with c*-MYC* have resulted in loss of the *c-MYC* promoter, either retaining or losing the G4 quadruplex (Raji and CA46, respectively; Fig. 6B) ^41^. DJ34 treatment down-regulated c-Myc in both cell lines, suggesting that DJ34 does not require the G4 quadruplex to inhibit *c-MYC* transcription (Fig. 6C). RT-qPCR experiments with CA46 cells confirmed that DJ34 treatment resulted in a strong reduction in *c-MYC* mRNA (Fig. 6D). Together, these data show that DJ34 inhibits transcription of *c-MYC* and that this occurs independently of the G4 quadruplex.

### DJ34 intercalates DNA and inhibits topoisomerase II but not topoisomerase I

DJ34 contains three planar aromatic rings, a feature often found in DNA-intercalating compounds. Furthermore, compounds that inhibit topoisomerases through DNA intercalation have been shown to reduce *c-MYC* transcription ^42,43^. We therefore tested whether DJ34 intercalates and/or inhibits topoisomerase (topo) I/II to block *c-MYC* transcription. First, we performed a topo I DNA unwinding/intercalation assay, in which compounds are incubated with either a supercoiled or a relaxed substrate plasmid in the presence of topo I (see Suppl. Fig. S7I for an overview). We found that incubation of DJ34 with topo I and either a supercoiled or a relaxed plasmid substrate exclusively produced supercoiled plasmids, showing that DJ34 does indeed intercalate, but does not inhibit topo I (Fig. 6E, lanes 5 and 12). DJ34 intercalated at lower concentrations than m-Amsacrine, a known DNA intercalator used to treat ALL (compare lanes 4-5 with lanes 6-7, and lanes 11-12 with lanes 13-14).

We then performed a topo II relaxation assay based on a supercoiled plasmid substrate that is relaxed by topo II, using m-Amsacrine and Doxorubicine (topo II poisons), ICRF193 (topo II catalytic inhibitor), and Irinotecan (topo I inhibitor with no effect on topo II) as controls. This revealed that DJ34 is a potent inhibitor of topo II, producing a strong supercoiled band on the DNA gel similar to that observed for ICRF193 (Fig. 6F, lanes 3 and 8, respectively). The DJ34 analog Acridine did not inhibit topo II, suggesting specificity of DJ34, and determining the relevant molecular features of DJ34 is the focus of an ongoing follow-up study. Collectively, these data demonstrate that DJ34 is a DNA intercalator and a strong inhibitor of topo II, but not topo I.

It has previously been shown that intron 1 of the *c-MYC* gene contains a sequence that reduces *c-MYC* transcription due to torsional tension that arises during transcription, and the activity of topo II is essential for relieving this tension and maintaining high levels of *c-MYC* transcription; indeed, expression of plasmid-encoded *c-MYC* cDNA lacking this intron does not require topo II activity ^42-45^. We analyzed c-Myc levels in the MIA-PaCa GPS-Myc cell line ^46^, which contains the endogenous, wild-type *c-MYC* alleles in addition to a randomly integrated *dsRED-IRES-GFP-c-MYC* construct (Fig. 6G). This *c-MYC* cDNA lacks all introns including the aforementioned transcription-reducing intron 1, which is targeted by topo II to promote high *c-MYC* expression ^42-45^. Interestingly, analysis of mRNA levels by RT-qPCR revealed that although endogenous *c-MYC* mRNA levels were strongly reduced, transcription of *GFP-c-MYC* was completely resistant to DJ34 (Fig. 6H). Taken together, these data are consistent with a model in which DNA intercalation by DJ34 poisons topo II to interfere with *c-MYC* transcription.

### DJ34 induces cell cycle arrest, cell differentiation and apoptosis

We next investigated the physiological consequences of DJ34 treatment. Flow cytometry analysis showed that DJ34 treatment resulted in G1/G0 cell cycle arrest and apoptosis (Fig. 7A), which was confirmed by PARP cleavage in multiple cancer cell lines (Fig. 7B and Suppl. Fig. S7J). In addition to causing apoptosis and cell cycle arrest, we found that DJ34 increased the expression levels of the differentiation marker CD25 while decreasing the levels of CD43, a marker of undifferentiated hematopoietic progenitors (Fig. 7C). DJ34 appeared to be more effective at inducing cell differentiation than imatinib, which is important, because c-Myc is known to mediate imatinib resistance by preventing imatinib-induced cell differentiation ^47^, and preventing differentiation is one of the mechanisms by which c-Myc promotes drug resistance in leukemia ^48^. Collectively, these data indicate that DJ34 triggers apoptosis, cell cycle arrest and differentiation.

**Figure 7.**
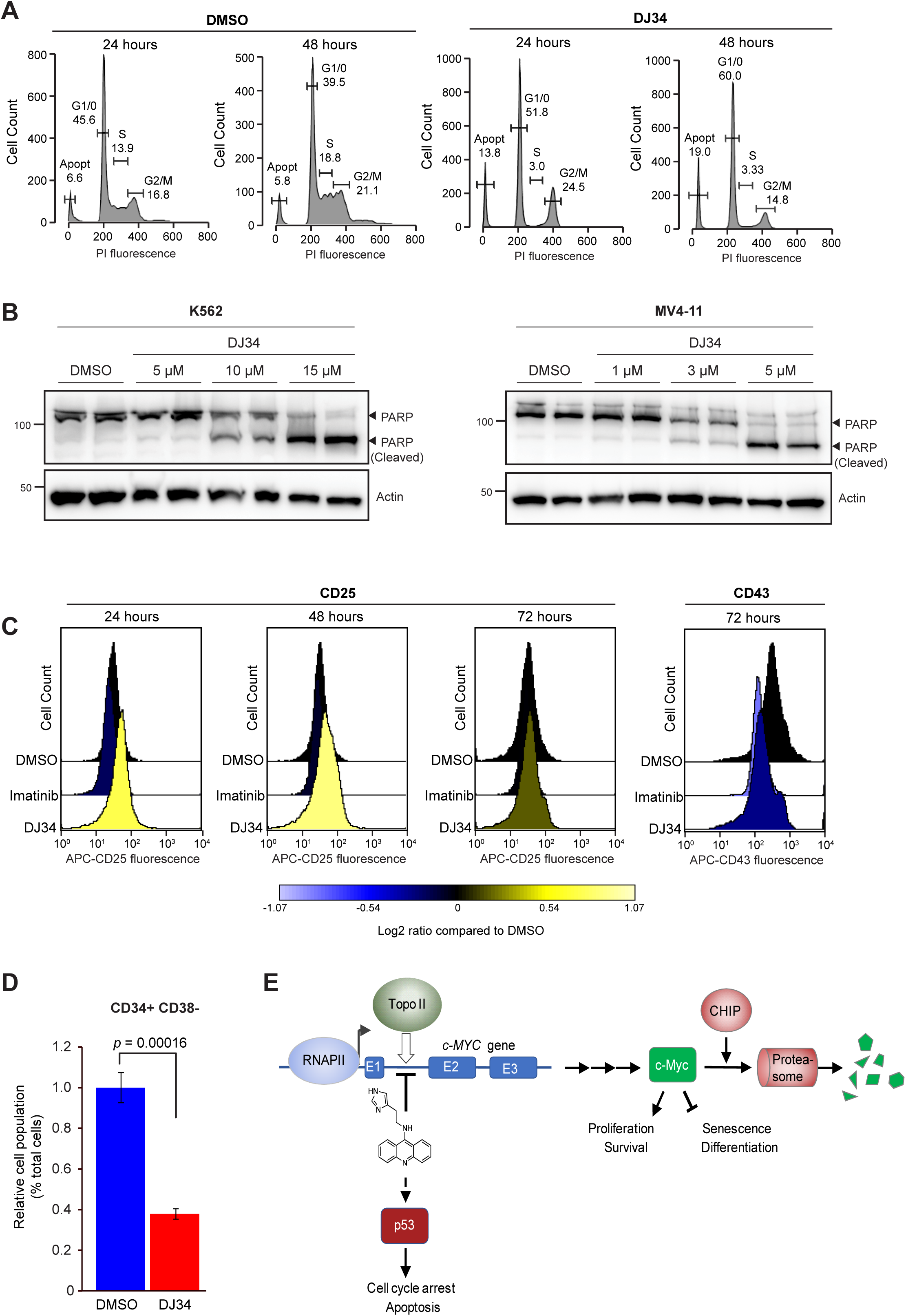
DJ34 induces cell cycle arrest, apoptosis and differentiation. (**A**) Cell cycle analysis of Ba/F3-BCR-Abl cells treated with DMSO or DJ34 for 24 hrs and 48 hrs. **(B)** Immunoblot analysis of K562 and MV4-11 cells treated with DMSO or DJ34 at the indicated concentrations for 48 hrs using antibodies against PARP and Actin. (**C**) Analysis by flow cytometry of the differentiation marker CD25 and the hematopoietic progenitor cell marker CD43 after treatment of Ba/F3-BCR-Abl cells with either imatinib or DJ34. 10,000 cells were counted for each treatment and timepoint. (**D**) CD34+CD38-LSCs are particularly sensitive to DJ34. Mononuclear cells were isolated from the bone marrow of a Ph+ CML patient and incubated for 24 hrs with 10 µM DJ34, after which the fraction of CD34+ CD38-cells in the total cell pool was analyzed by flow cytometry. Error bars, standard deviation. (**E**) Schematic overview of the findings of this study. DJ34 intercalates the DNA and inhibits topo II to activate p53 and to block transcription of *c-MYC*, which tentatively occurs in intron 1. c-Myc protein is then ubiquitinated and degraded by the proteasome through a pathway that requires CHIP.

### DJ34 has anti-LSC activity

LSCs mediate disease relapse, and novel therapy that eradicates LSCs is expected to provide durable remission ^49^. It was previously shown that inhibiting c-Myc and reactivating p53 is an effective approach to eradicate LSCs ^6^. Given that DJ34 has dual c-Myc and p53 targeting activity, we analyzed the effect of DJ34 on LSCs. We isolated bone marrow cells from a CML patient and treated the cells with either DMSO or with 10 μM DJ34. After 48 hrs we analyzed the relative number of LSCs in the total cell population by flow cytometry. Importantly, DJ34 significantly reduced the relative number of CD34+ CD38-LSCs in the total pool of bone-marrow-derived cancer cells (Fig. 7D), showing that LSCs are particularly sensitive to DJ34.

## Discussion

We performed a cell competition drug screen and identified several compounds that selectively inhibit BCR-Abl-expressing cells but not isogenic control cells. None of these compounds functioned as TKIs, suggesting they target parallel pathways that control proliferation and survival of transformed cells. Our findings support previous studies that showed that isogenic cell competition screens provide a powerful alternative approach to traditional cell viability-based drug screens, facilitating the identification of compounds that induce cell cycle arrest and cellular differentiation, as well as those that induce apoptosis. It has been argued that such phenotypic drug screens may help reduce the high attrition rates that have characterized drug development in the past decade ^50,51^.

We identified a compound that may provide an effective lead structure for therapy. A model of our findings is shown in Figure 7E. DJ34 is a DNA intercalator and inhibits topo II, resulting in activation of p53 and inhibition of *c-MYC* transcription. As demonstrated by RNAseq, DJ34 did not appear to broadly inhibit transcription and showed relative selectivity for *c-MYC*, as the main genes affected by DJ34 were *c-MYC* itself and c-Myc target genes. The exact DNA sequence recognized by DJ34 remains to be identified, and exactly how DJ34 activates p53 remains to be established, but it is likely similar to activation of p53 by other topo II poisons.

DJ34-induced depletion of c-Myc was independent of the phosphorylation status of c-Myc. This is important, because c-Myc-addicted tumors often stabilize c-Myc by interfering with its phosphorylation ^52^. Therefore, DJ34-based treatments may show clinical efficacy towards a broad range of c-Myc-addicted cancers, including cancers driven by oncogenic c-Myc mutants, such as the c-Myc-T58 and c-Myc-P57 mutations that are often found in Burkitt’s lymphoma and which are refractory to phospho-regulation ^53^. Furthermore, DJ34 overcame MLN4924-dependent stabilization of c-Myc, suggesting that DJ34 induces c-Myc depletion independently of SCF ubiquitin ligases. This provides a therapeutic advantage, because Cullins are often mutated and inactivated in cancer, including ALL ^37,54,55^. Indeed, loss of Fbxw7 has been shown to bolster leukemia-initiating stem cells in ALL by increasing c-Myc abundance, and inhibition of c-Myc resulted in remission in Fbxw7-mutant mouse models of ALL ^56^. We believe that DJ34, or its derivatives, may therefore be particularly effective in these forms of ALL. We are currently testing the efficacy of DJ34 in mouse models, and our preliminary data indicate that DJ34 has a favorable ADME/Tox profile and is well tolerated by animals, all of which will be reported elsewhere.

c-Myc induces stemness and blocks cellular senescence and differentiation in many forms of cancer and promotes LSC survival in leukemia. Brief or even partial suppression of c-Myc can result in acute and sustained tumor regression, and dual targeting of p53 and c-Myc is a particularly effective strategy to kill leukemic stem cells ^6^. However, development of drugs that inhibit c-Myc has progressed slowly, primarily because c-Myc is widely considered undruggable. Given that DJ34 has potent anti-c-Myc activity in a broad range of cancers, that it can simultaneously activate p53, and that LSCs are particularly sensitive to DJ34, we believe it forms an excellent starting point for further development of anti-c-Myc cancer therapy.

## Materials and Methods

### Cell culture

Suspension cells (Ba/F3, K-562, MEG-01, MV4-11, SD-1, CA46, Raji, Daudi, 697, RS411, and KU-812) were cultured with RPMI-1640 with 10% FBS and 1% penicillin/streptomycin (P/S). WT Ba/F3 cells also received 7.5 ng/ml IL-3. Adherent cells (MIA PaCa, MIA-PaCa - GPS-Myc, A549, H460, MCF7, U87, HCC-827, H1975, SKBR3, MRC5, A375, MEWO, SK-MEL28, WM1366, HCT116, MEFs and LN229) were cultured in DMEM with 10% FBS and 1% P/S. Primary patient cells were cultured with mononuclear cell basal medium (PromoCell, Heidelberg, Germany) supplemented with 10% SupplementMix (PromoCell, Heidelberg Germany).

### Cell competition-based drug screen

Compounds were diluted in 10% DMSO and dispensed at the Chemical Biology Screening Platform of the Nordic Centre for Molecular Medicine (Oslo, Norway) into individual wells of a 384-well plate using an Echo acoustic liquid dispenser (Labcyte, San Jose, CA, USA) such that, after addition of cells, the compound concentration was 5 µM. Drug libraries that were used were the Sigma LOPAC1280 library and the ChemBioNet drug library ^11^.

A total of 1500 cells were mixed at a BCR-Abl:WT cell ratio of 1.3:1 in 50 µl cell culture medium (containing IL3). This ratio accounted for the slower growth rate of cells transfected with BCR-Abl and ensured that the ratio after 72 hours treatment with DMSO was 1:1. In addition to library compounds, imatinib (5 µM) was used as a positive control in 16 wells of each 384-well plate. After 72 hours, the number of GFP-positive (BCR-Abl) and RFP-positive (WT) cells in each well was recorded by flow cytometry and used to calculate a cell ratio.

### Chemical synthesis

Synthesis of DJ34 and other compounds was performed by Hit2Lead, ChemBridge, Cambridge, UK. DJ34 is available upon request.

### RNA Sequencing

RNAseq was performed as previously described ^12^; see Supplementary Methods. Data were deposited at Gene Expression Omnibus (GEO; GSE100678).

### RT-qPCR

RT-qPCR experiments were performed as previously described ^13^; see Supplementary Methods.

### In vitro kinase assays

Kinase assays were performed using the Omnia Kinase Assay according to the manufacturer’s protocol (Life Technologies, Carlsbad, CA, USA); see Supplemental Methods.

### Proteomic and Phosphoproteomic analyses

MS experiments were performed as previously described ^14^, see Supplemental Methods. All MS data were deposited to the ProteomeXchange Consortium via PRIDE repository (PXD018812).

### Flow Cytometry and phosphoflow cytometry

Flow cytometry was performed as previously described ^15^. See Supplemental Methods.

### Differentiation analysis

Bcr/Abl-expressing Ba/F3 cells were treated with DMSO, 10 µM imatinib or 20 µM DJ34 for 24, 48 and 72 hours. Cells were fixed as described above (see cell cycle analysis), incubated with APC-conjugated mouse anti-CD-25 and CD-43 antibodies (BD Biosciences, San Jose CA, USA) and analyzed by flow cytometry for surface expression of CD25 and CD43.

### Apoptosis analysis

BCR-Abl-expressing Ba/F3 cells were treated with DMSO, 10 µM imatinib and 20 µM DJ34 for 24h, 48h and 72h. Samples were then analyzed by flow cytometry and cells undergoing apoptosis were quantified/gated according to changes in morphological and physical properties using forward and side scattered light.

### Western blotting

Western blotting was performed as previously described ^16^; see Supplemental Methods.

### ADME-PK analysis

ADME-PK analysis was outsourced to the public CRO Cyprotex (www.cyprotex.com).

### DNA intercalation and topoisomerase assays

DNA intercalation and topoisomerase I and II activity assays were performed using a DNA Unwinding Assay Kit and a Human Topoisomerase II Relaxation Assay Kit, respectively, as described in the user manual (Inspiralis, Norwich, UK).

### Patient information

Patient #10: Caucasian male, ALL; Patient #13: Caucasian male, Ph+ ALL; Patient #14: Caucasian female, AML; Patient #45: Caucasian male, CML.

### Ethical considerations

All samples were collected after obtaining written informed consent. The study was approved by the local ethical review board (REK2015/1012; Regional Ethics Committee South East, committee D) in accordance with the Declaration of Helsinki.

## Acknowledgements

The HCT116 WT and HCT116 FBXW7-/- cells were a kind gift from Dr. B. Vogelstein. GPS-Myc cells were a kind gift from Prof. Channing Der (University of North Carolina at Chapel Hill, NC, USA). We thank Dr. P. Chymkowitch for technical assistance, and Dr. G.A. De Souza for assistance with MS data analysis. The research leading to these results has received funding from the European Union Seventh Framework Programme (FP7-PEOPLE-2013-COFUND) under grant agreement n° 609020 - Scientia Fellows. This work was further supported by grants from the Norwegian Cancer Society (project numbers 3311782, 4487303, 144176, 6786517, 182524), the Norwegian Health Authority South-East (2014014, 2012012) and the Norwegian Research Council (267454). This work was partly supported by the Research Council of Norway through its Centres of Excellence funding scheme, project number 262652.

## Author contributions

Performed research: DST, JR, JS, LP, PAD, RSM, RC. Contributed vital reagents: YF, BTG, TGD. Analyzed data: JME, DST, JR, JS, LP, PAD, RSM, RC, SKS, JW, BTG. Wrote the paper: DST, JR, JME

## Supplementary Figure Legends

**Supplementary Figure S1**. (**A**) Overview of the name, known molecular targets and structure of the compounds that increase the relative competitiveness of BCR-Abl-expressing cells relative to isogenic control cells. The results obtained with the competition assay and cell viability assay are also shown. All values were normalized to DMSO. (**B**) Effect of the JAK2 inhibitors Ruxolitinib, S-Ruxolitinib and Tofacitinib on the viability of WT and BCR-Abl-expressing Ba/F3 cells. Cells were treated with the indicated doses of drugs for 72 hrs after which cell viability was assessed by CellTiter-Glo. (**C**) Relative viability of BCR-Abl-expressing or WT cells treated with the JAK2 kinase inhibitor AZD1480 combined with either DMSO or imatinib. Error bars: standard deviation.

**Supplementary Figure S2**. (**A**) The ratio of BCR-Abl-expressing cells over WT Ba/F3 cells following treatment for 72 hours with the topoisomerase II inhibitors ICRF-193 (upper x axis) and Etoposide, Doxorubicin and Camptothecin (lower x axis). Ratios were calculated using the cell competition-based assay (see Fig. 1). (**B-G**) Cell viability of BCR-Abl-expressing or WT Ba/F3 cells following treatment with the PDE inhibitors IBMX (*B*), Ibidulast (*C*), Rolipram (*D*) and Zardaverine (*E*); the adenylate cyclase activator forskolin (*F*), and the inactive forskolin analogue dideoxyforskolin for 72 hours (*G*). (**H, I**) Effect of the cAMP analogues 8Br-cAMP or 007 on the viability and competitiveness of BCR-Abl-expressing and WT Ba/F3 cells after 72 hours treatment.

**Supplementary Figure S3**. Relative kinase activity of purified recombinant Abl kinase domain following treatment with DJ1, DJ2, DJ3, DJ34 and DJ35. DMSO treatment was used as a negative control, and the Abl kinase inhibitor ponatinib was used as a positive control. All data were normalized to timepoint 0.

**Supplementary Figure S4. (A-D)** Cell viability of human leukemia cell lines following 72 hours treatment with DJ1, DJ2, DJ3, DJ12, DJ34, DJ35, 8Br-cAMP or forskolin. Cell lines used were the CML cell lines MEG-01 (*A*), KU-812 (*B*) and K562 (*C*), and SD-1 (Ph+ ALL) (*D*).

**Supplementary Figure S5**. (A) Phospho-flow cytometry analysis showing the effect of DJ34 or imatinib on the levels of phosphorylation sites known to be components of oncogenic signaling pathways. (**B**) Immunoblot analysis of BCR-Abl-expressing Ba/F3 cells treated with DJ34 or imatinib for 4 hrs. Membranes were probed with antibodies targeting phosphorylation sites that are known components of signaling pathways downstream of BCR-Abl.

**Supplementary Figure S6**. (**A**) Metascape analysis of the 1705 transcripts that increased at least 2-fold in abundance following RNA-seq analysis of DJ34-treated cells. (**B**) Total numbers of unique phosphopeptides identified by MS from cells treated with 10 µM imatinib or 20 µM DJ34.

**Supplementary Figure S7. (A)** Immunoblot analysis of BCR-Abl-expressing Ba/F3 cells treated with 10 µM DJ34 for 1, 2 or 4hrs after pre-treatment for 1hr with 10 µM MG132. (**B** and **C**) Immunoblot analyses of BCR-Abl-expressing Ba/F3 cells treated with DJ34 after 1 hr pre-treatment with either 500 nM okadaic acid (OA; *B*) or 10 µM MLN4924 (*C*). (**D**) DJ34-induced c-Myc depletion does not require SCF^FBW7^. WT HCT116 cells or isogenic FBXW7^-/-^ knock-out cells were mock-treated or treated with DJ34 for the indicated times, after which cell lysates were analyzed by immunoblotting with antibodies against c-Myc and Actin. Quantification of c-Myc bands across three technical repeat experiments is displayed below the immunoblotting images. (**E**) Immunoblot analysis showing the effect of DJ34 on cyclin E levels in three different cancer cell lines. (**F**) DJ34-induced c-Myc depletion is mediated by CHIP. WT MEFs or CHIP-/- MEFs were treated and analyzed as in (*D*). (**G**) DJ34 down-regulates c-Myc upstream of the ribosome. Cells were pretreated with 10 µg/ml cycloheximide for 10 mins, followed by treatment with DMSO or DJ34 for the indicated times, after which cell lysates were analyzed by immunoblotting with c-Myc and Actin antibodies. For comparison, cells were also treated with DJ34 alone (i.e. 10 mins DMSO pre-treatment followed by DJ34). (**H**) DJ34 does not accelerate α-amanitin-induced loss of c-Myc. Cells were treated with 50 µM α-amanitin and 10 µM DJ34 as indicated, after which cell lysates were analyzed by western blotting with c-Myc and Actin antibodies. (**I**) Schematic overview of the topo I DNA unwinding/intercalation assay (with both supercoiled and relaxed DNA plasmids) outlining the possible results for a compound with different DNA intercalating and topo I inhibitory properties. Lanes represent the different properties in relation to whether or not a compound can intercalate DNA or inhibit topo I, respectively, which is further summarized in the table underneath the two panels. In brief, in this hypothetical experiment a compound that neither intercalates nor inhibits topo I will produce a relaxed plasmid for both substrates (compound A, lane 3 in both panels), whereas a compound that does intercalate but does not inhibit topo I will produce supercoiled plasmids, regardless of the substrate used (compound B, lane 4 in both panels). Finally, a compound that both intercalates DNA and inhibits topo I will produce a supercoiled plasmid from a supercoiled substrate, and a relaxed plasmid from a relaxed substrate (compound C, lane 5 in both panels). (**J**) Immunoblot analysis of RS4-11 and LN229 cells treated with DMSO or DJ34 at the indicated concentrations for 48 hrs. Cell lysates were analyzed with antibodies against PARP and Actin.

**Supplemental Figure S8**. Uncropped western blots shown in Figure 4.

**Supplemental Figure S9**. Uncropped western blots shown in Figure.

**Supplemental Figure S10**. Uncropped western blots shown in Figure 7.

**Supplemental Figure S11**. Uncropped western blots shown in Suppl. Figure S5.

**Supplemental Figure S12**. Uncropped western blots shown in Suppl. Figure S7.

## References

1 Rowley, J. D. Genetics. A story of swapped ends. Science 340, 1412–1413, doi: 10.1126/science.1241318 (2013).

2 Roberts, K. G. & Mullighan, C. G. Genomics in acute lymphoblastic leukaemia: insights and treatment implications. Nature Reviews Clinical Oncology 12, 344–357, doi: 10.1038/nrclinonc.2015.38 (2015).

3 Fielding, A. K. et al. UKALLXII/ECOG2993: addition of imatinib to a standard treatment regimen enhances long-term outcomes in Philadelphia positive acute lymphoblastic leukemia. Blood 123, 843–850, doi: 10.1182/blood-2013-09-529008 (2014).

4 Cilloni, D. & Saglio, G. Molecular pathways: BCR-ABL. Clin Cancer Res 18, 930–937, doi: 10.1158/1078-0432.ccr-10-1613 (2012).

5 Sawyers, C. L., Callahan, W. & Witte, O. N. Dominant negative MYC blocks transformation by ABL oncogenes. Cell 70, 901–910, doi: 10.1016/0092-8674(92)90241-4 (1992).

6 Abraham, S. A. et al. Dual targeting of p53 and c-MYC selectively eliminates leukaemic stem cells. Nature 534, 341–346, doi: 10.1038/nature18288 (2016).

7 Gabay, M., Li, Y. & Felsher, D. W. MYC activation is a hallmark of cancer initiation and maintenance. Cold Spring Harb Perspect Med 4, doi: 10.1101/cshperspect.a014241 (2014).

8 Bonnet, D. & Dick, J. E. Human acute myeloid leukemia is organized as a hierarchy that originates from a primitive hematopoietic cell. Nat Med 3, 730–737, doi: 10.1038/nm0797-730 (1997).

9 Corbin, A. S. et al. Human chronic myeloid leukemia stem cells are insensitive to imatinib despite inhibition of BCR-ABL activity. J Clin Invest 121, 396–409, doi: 10.1172/jci35721 (2011).

10 Schepers, K., Campbell, T. B. & Passegue, E. Normal and leukemic stem cell niches: insights and therapeutic opportunities. Cell Stem Cell 16, 254–267, doi: 10.1016/j.stem.2015.02.014 (2015).

11 Lisurek, M. et al. Design of chemical libraries with potentially bioactive molecules applying a maximum common substructure concept. Molecular Diversity 14, 401–408, doi: 10.1007/s11030-009-9187-z (2010).

12 Chymkowitch, P. et al. Sumoylation of Rap1 mediates the recruitment of TFIID to promote transcription of ribosomal protein genes. Genome Res 25, 897–906, doi: 10.1101/gr.185793.114 (2015).

13 Herrera, M. C. et al. Cdk1 gates cell cycle-dependent tRNA synthesis by regulating RNA polymerase III activity. Nucleic Acids Res 46, 11698–11711, doi: 10.1093/nar/gky846 (2018).

14 Robertson, J. et al. Defining the phospho-adhesome through the phosphoproteomic analysis of integrin signalling. Nat Commun 6, 6265, doi: 10.1038/ncomms7265 (2015).

15 Irish, J. M. et al. Single cell profiling of potentiated phospho-protein networks in cancer cells. Cell 118, 217–228, doi: 10.1016/j.cell.2004.06.028 (2004).

16 Enserink, J. M. et al. A novel Epac-specific cAMP analogue demonstrates independent regulation of Rap1 and ERK. Nat Cell Biol 4, 901–906, doi: 10.1038/ncb874 (2002).

17 Asmussen, J. et al. MEK-Dependent Negative Feedback Underlies BCR–ABL-Mediated Oncogene Addiction. Cancer Discovery 4, 200, doi: 10.1158/2159-8290.CD-13-0235 (2014).

18 Modi, H. et al. Role of BCR/ABL gene-expression levels in determining the phenotype and imatinib sensitivity of transformed human hematopoietic cells. Blood 109, 5411–5421, doi: 10.1182/blood-2006-06-032490 (2007).

19 Hantschel, O. et al. BCR-ABL uncouples canonical JAK2-STAT5 signaling in chronic myeloid leukemia. Nat Chem Biol 8, 285–293, doi: 10.1038/nchembio.775 (2012).

20 Boehrer, S. et al. Erlotinib exhibits antineoplastic off-target effects in AML and MDS: a preclinical study. Blood 111, 2170–2180, doi: 10.1182/blood-2007-07-100362 (2008).

21 Bogeso, K. P., Liljefors, T., Arnt, J., Hyttel, J. & Pedersen, H. Octoclothepin enantiomers. A reinvestigation of their biochemical and pharmacological activity in relation to a new receptor-interaction model for dopamine D-2 receptor antagonists. J Med Chem 34, 2023–2030, doi: 10.1021/jm00111a015 (1991).

22 Fajardo, A. M., Piazza, G. A. & Tinsley, H. N. The role of cyclic nucleotide signaling pathways in cancer: targets for prevention and treatment. Cancers (Basel) 6, 436–458, doi: 10.3390/cancers6010436 (2014).

23 Daina, A., Michielin, O. & Zoete, V. SwissADME: a free web tool to evaluate pharmacokinetics, drug-likeness and medicinal chemistry friendliness of small molecules. Sci Rep 7, 42717, doi: 10.1038/srep42717 (2017).

24 Zeller, K. I., Jegga, A. G., Aronow, B. J., O’Donnell, K. A. & Dang, C. V. An integrated database of genes responsive to the Myc oncogenic transcription factor: identification of direct genomic targets. Genome Biology 4, R69, doi: 10.1186/gb-2003-4-10-r69 (2003).

25 Li, Z. et al. A global transcriptional regulatory role for c-Myc in Burkitt’s lymphoma cells. Proc Natl Acad Sci U S A 100, 8164–8169, doi: 10.1073/pnas.1332764100 (2003).

26 Vafa, O. et al. c-Myc can induce DNA damage, increase reactive oxygen species, and mitigate p53 function: a mechanism for oncogene-induced genetic instability. Mol Cell 9, 1031–1044, doi: 10.1016/s1097-2765(02)00520-8 (2002).

27 Zheng, H. et al. Pten and p53 converge on c-Myc to control differentiation, self-renewal, and transformation of normal and neoplastic stem cells in glioblastoma. Cold Spring Harb Symp Quant Biol 73, 427–437, doi: 10.1101/sqb.2008.73.047 (2008).

28 Arango, D., Corner, G. A., Wadler, S., Catalano, P. J. & Augenlicht, L. H. c-myc/p53 interaction determines sensitivity of human colon carcinoma cells to 5-fluorouracil in vitro and in vivo. Cancer Res 61, 4910–4915 (2001).

29 Bouaoun, L. et al. TP53 Variations in Human Cancers: New Lessons from the IARC TP53 Database and Genomics Data. Hum Mutat 37, 865–876, doi: 10.1002/humu.23035 (2016).

30 Law, J. C., Ritke, M. K., Yalowich, J. C., Leder, G. H. & Ferrell, R. E. Mutational inactivation of the p53 gene in the human erythroid leukemic K562 cell line. Leuk Res 17, 1045–1050, doi: 10.1016/0145-2126(93)90161-d (1993).

31 Wang, Z. et al. Critical roles of p53 in epithelial-mesenchymal transition and metastasis of hepatocellular carcinoma cells. PLoS One 8, e72846, doi: 10.1371/journal.pone.0072846 (2013).

32 Makinoshima, H. et al. Signaling through the Phosphatidylinositol 3-Kinase (PI3K)/Mammalian Target of Rapamycin (mTOR) Axis Is Responsible for Aerobic Glycolysis mediated by Glucose Transporter in Epidermal Growth Factor Receptor (EGFR)-mutated Lung Adenocarcinoma. J Biol Chem 290, 17495–17504, doi: 10.1074/jbc.M115.660498 (2015).

33 Farrell, A. S. & Sears, R. C. MYC degradation. Cold Spring Harbor perspectives in medicine 4, a014365, doi: 10.1101/cshperspect.a014365 (2014).

34 Kuo, Y. C. et al. Regulation of phosphorylation of Thr-308 of Akt, cell proliferation, and survival by the B55alpha regulatory subunit targeting of the protein phosphatase 2A holoenzyme to Akt. J Biol Chem 283, 1882–1892, doi: 10.1074/jbc.M709585200 (2008).

35 Osaka, F. et al. A new NEDD8-ligating system for cullin-4A. Genes Dev 12, 2263–2268, doi: 10.1101/gad.12.15.2263 (1998).

36 Soucy, T. A. et al. An inhibitor of NEDD8-activating enzyme as a new approach to treat cancer. Nature 458, 732–736, doi: 10.1038/nature07884 (2009).

37 Akhoondi, S. et al. FBXW7/hCDC4 is a general tumor suppressor in human cancer. Cancer Res 67, 9006–9012, doi: 10.1158/0008-5472.can-07-1320 (2007).

38 Koepp, D. M. et al. Phosphorylation-dependent ubiquitination of cyclin E by the SCFFbw7 ubiquitin ligase. Science 294, 173–177, doi: 10.1126/science.1065203 (2001).

39 Kaplan, C. D., Larsson, K. M. & Kornberg, R. D. The RNA polymerase II trigger loop functions in substrate selection and is directly targeted by alpha-amanitin. Mol Cell 30, 547–556, doi: 10.1016/j.molcel.2008.04.023 (2008).

40 Dutta, D. et al. Cell penetrating thiazole peptides inhibit c-MYC expression via site-specific targeting of c-MYC G-quadruplex. Nucleic Acids Res 46, 5355–5365, doi: 10.1093/nar/gky385 (2018).

41 Brown, R. V., Danford, F. L., Gokhale, V., Hurley, L. H. & Brooks, T. A. Demonstration that drug-targeted down-regulation of MYC in non-Hodgkins lymphoma is directly mediated through the promoter G-quadruplex. J Biol Chem 286, 41018–41027, doi: 10.1074/jbc.M111.274720 (2011).

42 Collins, I., Weber, A. & Levens, D. Transcriptional consequences of topoisomerase inhibition. Mol Cell Biol 21, 8437–8451, doi: 10.1128/mcb.21.24.8437-8451.2001 (2001).

43 Bunch, R. T. et al. Influence of amsacrine (m-AMSA) on bulk and gene-specific DNA damage and c-myc expression in MCF-7 breast tumor cells. Biochem Pharmacol 47, 317–329, doi: 10.1016/0006-2952(94)90023-x (1994).

44 Chung, J., Sinn, E., Reed, R. R. & Leder, P. Trans-acting elements modulate expression of the human c-myc gene in Burkitt lymphoma cells. Proc Natl Acad Sci U S A 83, 7918–7922, doi: 10.1073/pnas.83.20.7918 (1986).

45 Spencer, C. A. & Groudine, M. Control of c-myc regulation in normal and neoplastic cells. Adv Cancer Res 56, 1–48, doi: 10.1016/s0065-230x(08)60476-5 (1991).

46 Vaseva, A. V. et al. KRAS Suppression-Induced Degradation of MYC Is Antagonized by a MEK5-ERK5 Compensatory Mechanism. Cancer Cell 34, 807–822 e807, doi: 10.1016/j.ccell.2018.10.001 (2018).

47 Gomez-Casares, M. T. et al. MYC antagonizes the differentiation induced by imatinib in chronic myeloid leukemia cells through downregulation of p27(KIP1.). Oncogene 32, 2239–2246, doi: 10.1038/onc.2012.246 (2013).

48 Pan, X.-N. et al. Inhibition of c-Myc overcomes cytotoxic drug resistance in acute myeloid leukemia cells by promoting differentiation. PloS one 9, e105381–e105381, doi: 10.1371/journal.pone.0105381 (2014).

49 Dick, J. E. Stem cell concepts renew cancer research. Blood 112, 4793–4807, doi: 10.1182/blood-2008-08-077941 (2008).

50 Swinney, D. C. & Anthony, J. How were new medicines discovered? Nat Rev Drug Discov 10, 507–519, doi: 10.1038/nrd3480 (2011).

51 Moffat, J. G., Vincent, F., Lee, J. A., Eder, J. & Prunotto, M. Opportunities and challenges in phenotypic drug discovery: an industry perspective. Nat Rev Drug Discov 16, 531–543, doi: 10.1038/nrd.2017.111 (2017).

52 Kalkat, M. et al. MYC Deregulation in Primary Human Cancers. Genes 8, 151, doi: 10.3390/genes8060151 (2017).

53 Bahram, F., von der Lehr, N., Cetinkaya, C. & Larsson, L. G. c-Myc hot spot mutations in lymphomas result in inefficient ubiquitination and decreased proteasome-mediated turnover. Blood 95, 2104–2110 (2000).

54 Malyukova, A. et al. The tumor suppressor gene hCDC4 is frequently mutated in human T-cell acute lymphoblastic leukemia with functional consequences for Notch signaling. Cancer Res 67, 5611–5616, doi: 10.1158/0008-5472.can-06-4381 (2007).

55 Heo, J., Eki, R. & Abbas, T. Deregulation of F-box proteins and its consequence on cancer development, progression and metastasis. Semin Cancer Biol 36, 33–51, doi: 10.1016/j.semcancer.2015.09.015 (2016).

56 King, B. et al. The ubiquitin ligase FBXW7 modulates leukemia-initiating cell activity by regulating MYC stability. Cell 153, 1552–1566, doi: 10.1016/j.cell.2013.05.041 (2013).

