## Supplemental material for "A cell competition-based drug screen identifies a novel compound that induces dual c-Myc depletion and p53 activation"

#### Supplemental Materials and Methods

##### RNA sequencing

BCR-Abl-expressing Ba/F3 cells were treated with DMSO or 20 μM DJ34 in triplicate for 4 hrs. Total RNA was extracted using the RNeasy mini kit according to the manufacturer's protocol (Qiagen, Manchester, UK). Purity of the RNA was confirmed by bioanalyzer, and RNA transcriptome library construction and Illumina Hiseq 4000 was performed at BGI Tech solutions (Hong Kong). RNA sequencing (RNA Seq) analysis was performed by Hemispherian RS (Oslo, Norway).

Prior to follow-up analyses, a value of 0.1 was added to the mean FPKM RNA Seq readings of all significant genes, so that genes with a zero value in one condition were still assigned a fold-change value when DMSO and DJ34 were compared. Genes with FPKM RNA Seq readings lower than one in both conditions were removed. Integrative genomics viewer (IGV) plots showing RNA Seq data for specific genes were generated using the IGV visualization tool as described by Robinson et al[1]. Gene set enrichment analysis (GSEA) was performed with GSEA software [2] using all genes that were significantly altered by DJ34 treatment. Metascape analysis (http://metascape.org [3]) was performed using only genes that were more than 2-fold up-regulated by DJ34 treatment. Enrichment cut-offs for Metascape were p-value<0.01, minimum gene count 3 and enrichment factor >1.5.

##### RT-qPCR

BCR-Abl expressing Ba/F3 cells were treated with DMSO or 20μM DJ34 for 4 hrs and total RNA were extracted as described for RNA sequencing. For each treatment, 100 ng of total RNA was used to synthesize cDNA using the QuantiTect reverse transcription kit according to the manufacturers protocol (Qiagen). RT-qPCR was performed using 2x fast syber green master mix (Foster City CA, USA), 0.2 μM of forward and reverse primers for the MYC target genes CDK4, EIf4e, and GAD45. Primers used for qPCR were 5'TGTTTGAGCATATAGACCAG3' forward and 5'AATCCCAACAACTTCTATTG3' reverse for CDK4, 5'CTACTGATGGACACTTCTAC3' forward and 5’ACTTAGAGATCAATCGAAGG3’ reverse for EIF4E, 5’GAGAACGACATCAACATC3’ forward and 5’CTCGGACAAGGTCCA3’ reverse for GAD45, 5’TCGGGTAGTGGAAAACCAGC3’ forward and 5’TTCCTGTTGGTGAAGCTAACGTT3’ reverse for MYC (exon1), 5’Ccgaccagctggag3’ forward and 5’CAgcttctctgagacgagct3’ reverse for MYC (exon2), 5’CAGCACGACTTCTTCAAGTCCG3’ forward and 5’GTAGTTGTACTCCAGCTTGTGCC3’ reverse primers for MYC (EGFP-exon2), and 5’TCGTCCCGTAGACAAAATGGT3’ forward and 5’CGCCCAATACGGCCAAA3’ reverse for GAPDH.

**Proteomics**

BCR-Abl-expressing Ba/F3 cells were treated with DMSO, 10 μM imatinib or 20 μM DJ34 for 4 hrs. Cells were lysed using a Triton-based lysis buffer (150 mM NaCl, 25 mM Tris pH 7.4, 1% (w/v) Titon-X-100) containing a protease inhibitor cocktail (Calbiochem Set 1; Merck Millipore) and a phosphatase inhibitor cocktail (Thermo Fisher Scientific). Samples equating to 1 mg of protein were reduced using dithiotheitol (DTT), precipitated using -20°C acetone and proteolytically digested using 20 μg trypsin (Promega, Madison, WI, USA).

For the proteomic element of the workflow, approximately 50 μg of peptides was de-salted using $C_{18}$ StageTips and analyzed by mass spectrometry (MS) to identify protein components.

The remaining peptides (approximately 950 μg) were processed for identification of phosphopeptides. Samples were de-salted using Oasis HLB sample extraction columns (Waters, Manchester, UK) and enriched for phosphopeptides using titanium dioxide enrichment beads (see Robertson et al [4] for details). Enriched fractions were de-salted twice using reverse-phase ZipTips containing $C_{18}$ media (Merck Millipore), and analyzed by MS to identify phosphopeptides.

Tandem MS analysis (LC-MS/MS) of all samples was performed in triplicate using an Easy nLC1000 liquid chromatography (LC) system (Thermo Electron, Bremen, Germany) coupled to a QExactive Plus Hybrid Quadrupole-Orbitrap mass spectrometer (Thermo Electron) with a nanoelectrospray ion source (EasySpray, Thermo Electron). The LC separation of peptides was performed using an EasySpray C18 analytical column (2 µm particle size, 100 Å, 75 µm inner diameter; Thermo Fisher Scientific). Peptides were separated over a 120 min solvent gradient from 2% to 30% (v/v) ACN in 0.1% (v/v) FA, after which the column was washed using 90% (v/v) ACN in 0.1% (v/v) FA for 20 min (flow rate 0.3 µL/min). All LC-MS/MS analyses were operated in data-dependent mode where the most intense peptides were automatically selected for fragmentation by high energy collision-induced dissociation.

Raw files from MS analyses were submitted to MaxQuant software [5] for peptide/protein identification using the Uniprot mouse database. MaxQuant output files (proteinGroups.txt for proteomic data and STY(sites).txt for phosphoproteomic data) were loaded into the Perseus software [6]. Identifications from potential contaminants and reversed sequences were removed and intensities were transformed to log2. Identified phosphorylation sites were filtered only for those that were confidently localized (class I, localization probability ≥ 0.75). All zero intensity values were replaced using noise values of the normal distribution of each sample. For proteomic data, protein abundances were compared using LFQ intensity values and a two-sample Student's T-test (permutation-based FDR correction (250 randomizations), FDR cut-off: 0.05, S0: 0.1). For phosphoproteomic data, phosphosite abundances were compared using intensity values and a two-sample Student's T-test (p value cut off: 0.05). Only phosphosites that were significantly increased or decreased by DJ34 in both of two biological repeat experiments were considered for follow up. For display of phosphoproteomic data, volcano plots were generated in Perseus, with phosphosites labelled as significantly decreased according to the Student's T-test. Lists of all identified proteins and phosphosites, as well as those that were significantly affected by DJ34 or imatinib treatment, can be found in Supplementary Tables S4 and S5.

**Flow Cytometry and phosphoflow cytometry**

BCR-Abl-expressing Ba/F3 cells were treated with DMSO or 20 µM DJ34 for 24 and 48 hours. Cells were washed in cold PBS and fixed in cold 70% (v/v) ethanol for 2 hours. Subsequently, fixed cells were stained with a solution containing 0.1% (w/v) Triton-X-100, 10 µg/ml propidium iodide and 100 µg/ml RNase A in PBS at 37°C for 10 min and analyzed by flow cytometry. For LSC analysis, primary patient cells were recovered by over-night culturing, then cells were treated with DMSO or 10 µM DJ 34 for 24 hours. Cells were washed in cold PBS and blocked with 0.5% BSA for 45 minutes. Cells were then stained with FITC conjugated anti-CD38 and PE conjugated anti-CD34 antibodies (Abcam, Cambridge, UK) for 45 minutes and analyzed by flow cytometry.

**In vitro kinase assays**

10 µg/ml of purified GST-Abl kinase domain was incubated with 10 µM of the Omnia Tyrosine Peptide Y6, 0.2 mM DTT, 10 µM of the compounds of interest, and 50 µM ATP. Fluorescence intensity was measured using the Envision 2104 Multilabel Reader (Perkin Elmer; Waltham, MA, USA). Fluorescence intensities were normalized to the first time-point measured.

**Western blotting**

Cells were treated with compounds or, as a control, DMSO. Unless stated otherwise, DJ34 was used at 20 µM and imatinib was used at 10 µM, and the incubation time was 4 hours. Cells were washed with PBS and lysed using Laemmli buffer or RIPA buffer (150 mM NaCl, 50 mM Tris-Hcl pH 8.0, 1% (w/v) Triton-X-100, 0.5% (w/v) sodium deoxycholate, 0.1% (w/v) SDS) containing a protease inhibitor cocktail (Calbiochem Set 1; Merck Millipore) and a phosphatase inhibitor cocktail (Thermo Fisher Scientific, Waltham, MA, USA). Lysates were centrifuged to remove insoluble material (22,000 *g*, 15 min, 4°C). Proteins were separated by SDS-PAGE and transferred to a nitrocellulose membrane (Bio-Rad, Hercules, CA, USA). Membranes were blocked using blocking buffer (5% (w/v) skim milk (Sigma-Aldrich) in TBS containing 0.1% (v/v) Tween-20 (TBS-T)) and probed overnight at 4°C with primary antibodies incubated in blocking buffer. Membranes were washed for

30 mins using TBS-T, and then incubated with the appropriate horseradish peroxidase (HRP)-conjugated secondary antibodies (GE Healthcare, USA) in blocking buffer for 60 min. Membranes were washed for 30 min using TBS, incubated with enhanced chemiluminiscent substrate for HRP (Thermo Fisher Scientific) for 5 min and scanned for visualization. For a list of antibodies see Supplemental Table S1.

**Plasmids**

Commercially available plasmids used were pcDNA3::p210-BCR-Abl, which was a gift from Warren Pear (Addgene plasmid # 27481, Cambridge MA, USA); GFP expression, pRNAT-H1.1/Hygro plasmid from Genscript (Piscataway NJ, USA); RFP expression, pmCherry-N1 from Clontech (Mountain View CA, USA). The plasmid used for expression of recombinant Abl kinase domain (c-Abl amino acids 220-498) for in vitro kinase assays was a kind gift from Michael W. Deininger (University of Utah Huntsman Cancer Institute, UT, USA). All transfections were performed using the Amaxa Biosystems Nucleofector II (Lonza, Cologne, Germany).

**References**


1. Robinson JT, Thorvaldsdottir H, Winckler W, Guttman M, Lander ES, Getz G, Mesirov JP: **Integrative genomics viewer.** *Nat Biotechnol* 2011, **29:**24-26.
2. Subramanian A, Kuehn H, Gould J, Tamayo P, Mesirov JP: **GSEA-P: a desktop application for Gene Set Enrichment Analysis.** *Bioinformatics* 2007, **23:**3251-3253.
3. Tripathi S, Pohl MO, Zhou Y, Rodriguez-Frandsen A, Wang G, Stein DA, Moulton HM, DeJesus P, Che J, Mulder LC, et al: **Meta- and Orthogonal Integration of Influenza "OMICs" Data Defines a Role for UBR4 in Virus Budding.** *Cell Host Microbe* 2015, **18:**723-735.
4. Robertson J, Jacquemet G, Byron A, Jones MC, Warwood S, Selley JN, Knight D, Humphries JD, Humphries MJ: **Defining the phospho-adhesome through the phosphoproteomic analysis of integrin signalling.** *Nat Commun* 2015, **6:**6265.
5. Cox J, Mann M: **MaxQuant enables high peptide identification rates, individualized p.p.b.-range mass accuracies and proteome-wide protein quantification.** *Nat Biotechnol* 2008, **26:**1367-1372.
6. Tyanova S, Temu T, Sinitcyn P, Carlson A, Hein MY, Geiger T, Mann M, Cox J: **The Perseus computational platform for comprehensive analysis of (prote)omics data.** *Nat Methods* 2016, **13:**731-740.

### Supplementary Figure S1

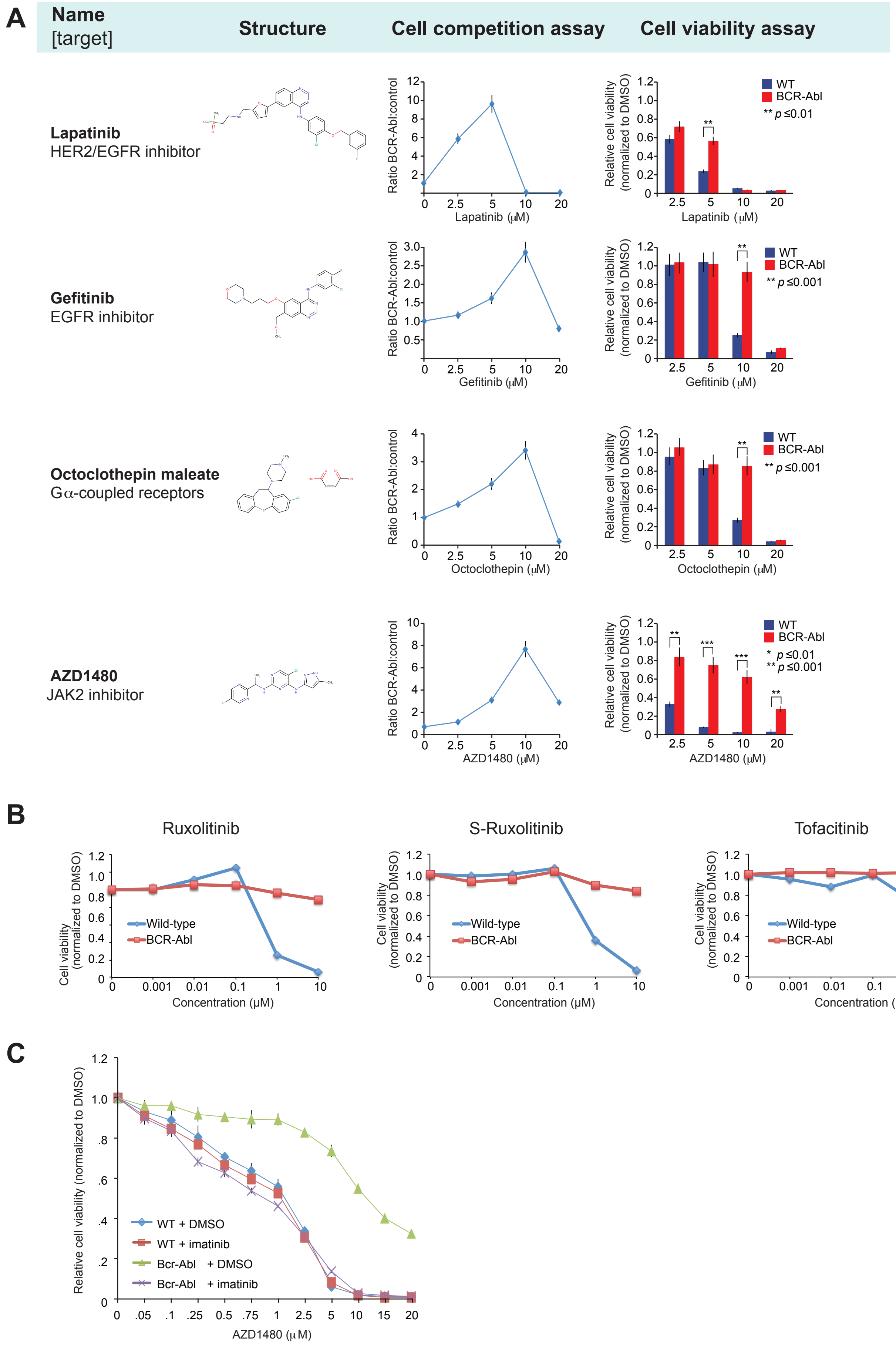

### Supplementary Figure S2

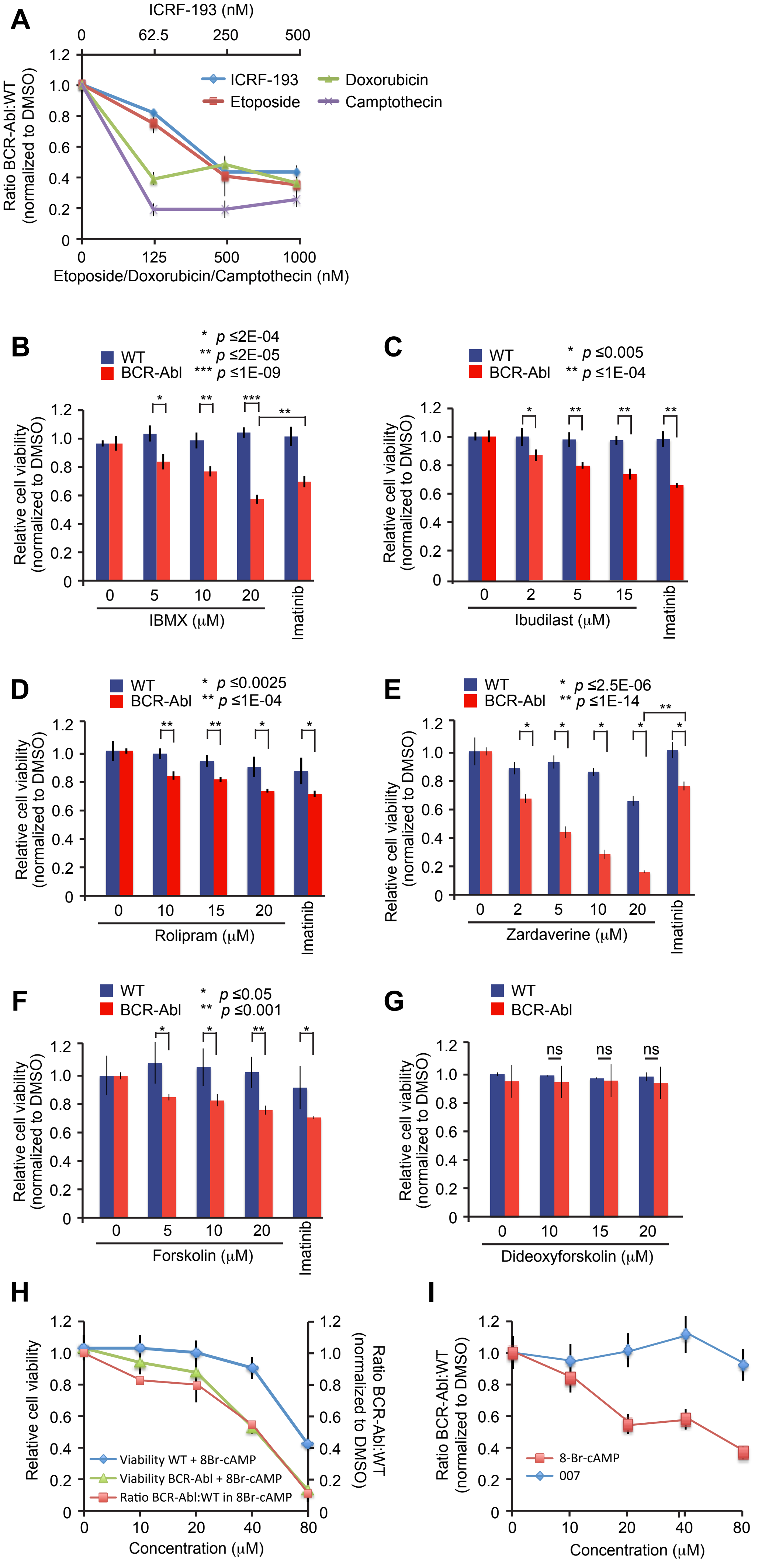

### Supplementary Figure S3

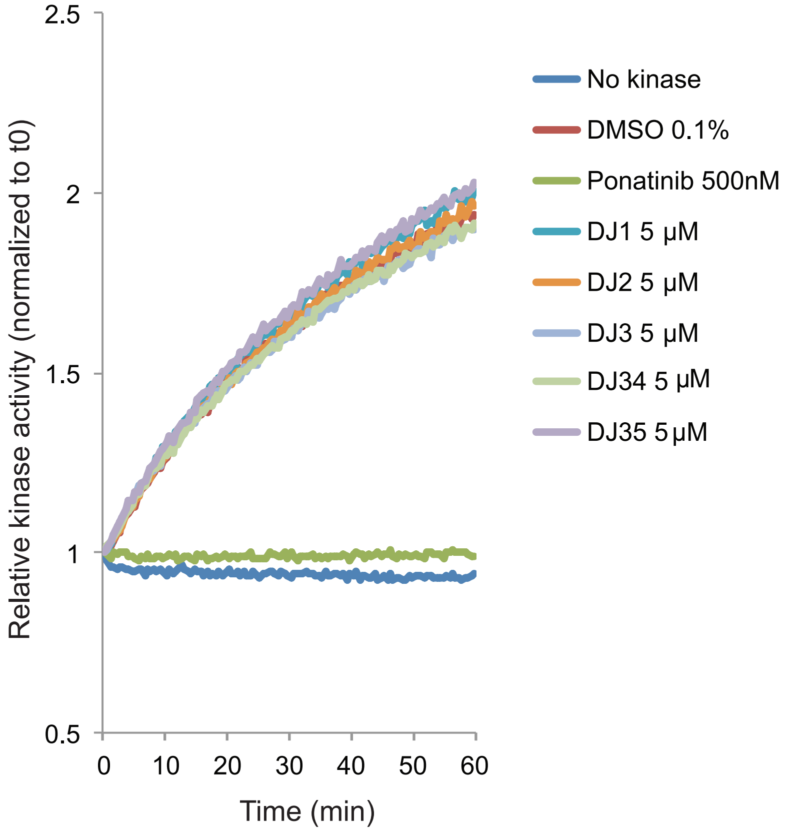

### Supplementary Figure S4

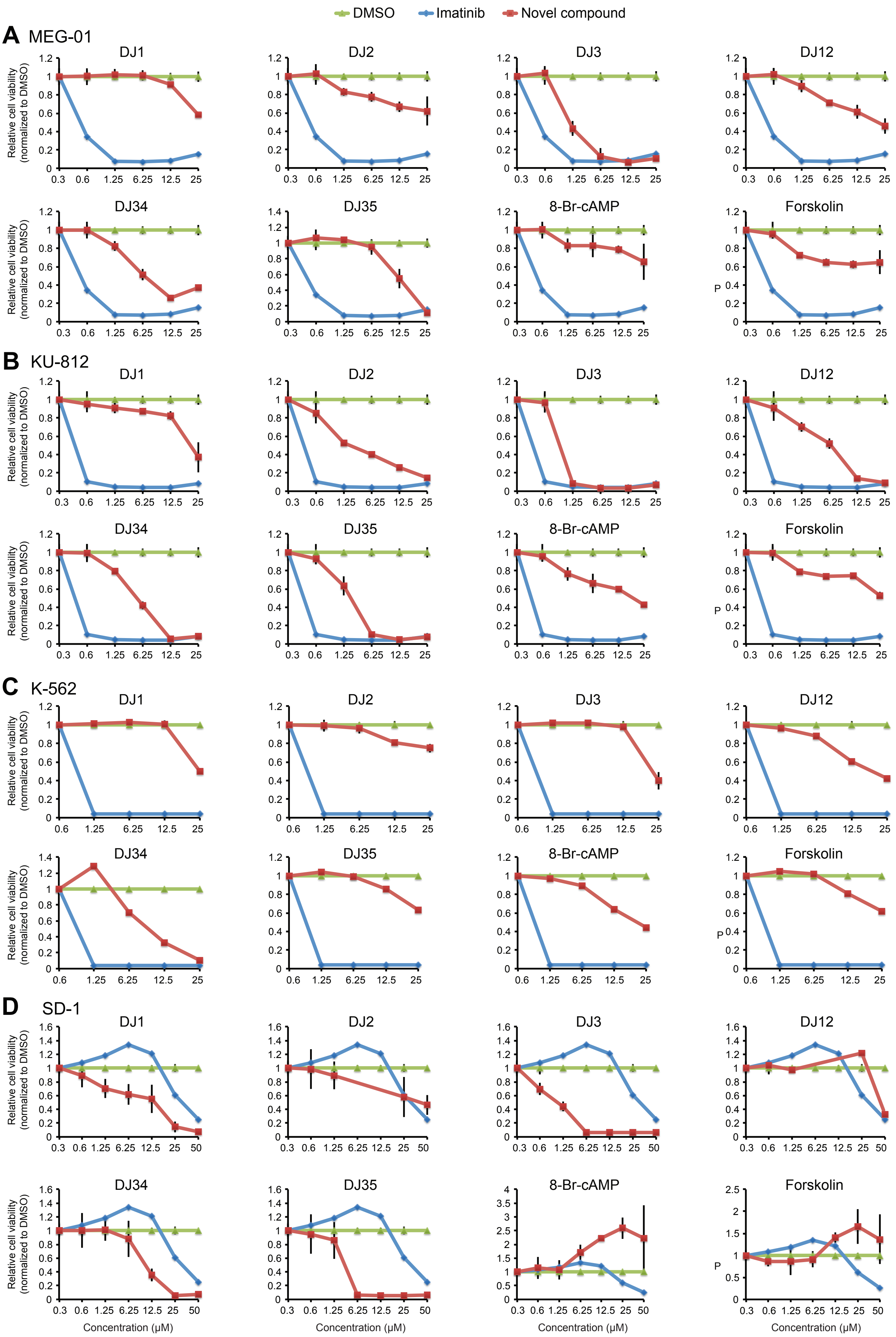

### Supplementary Figure S5

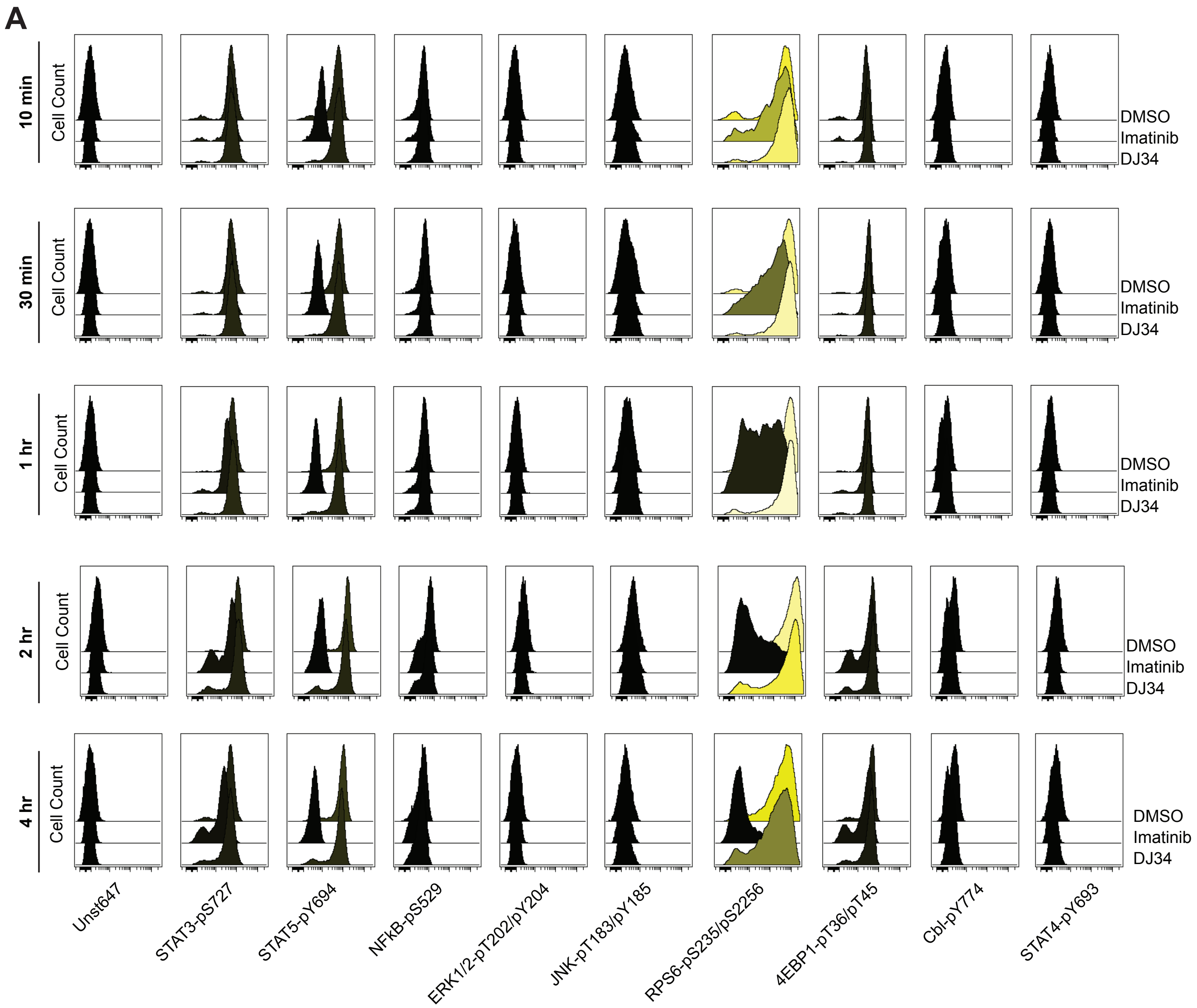


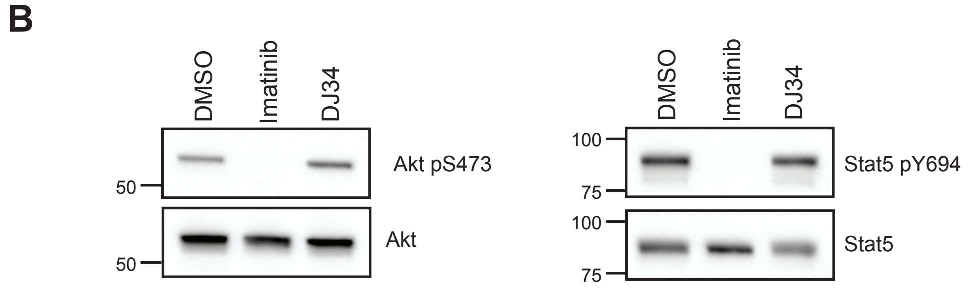

### Supplementary Figure S6

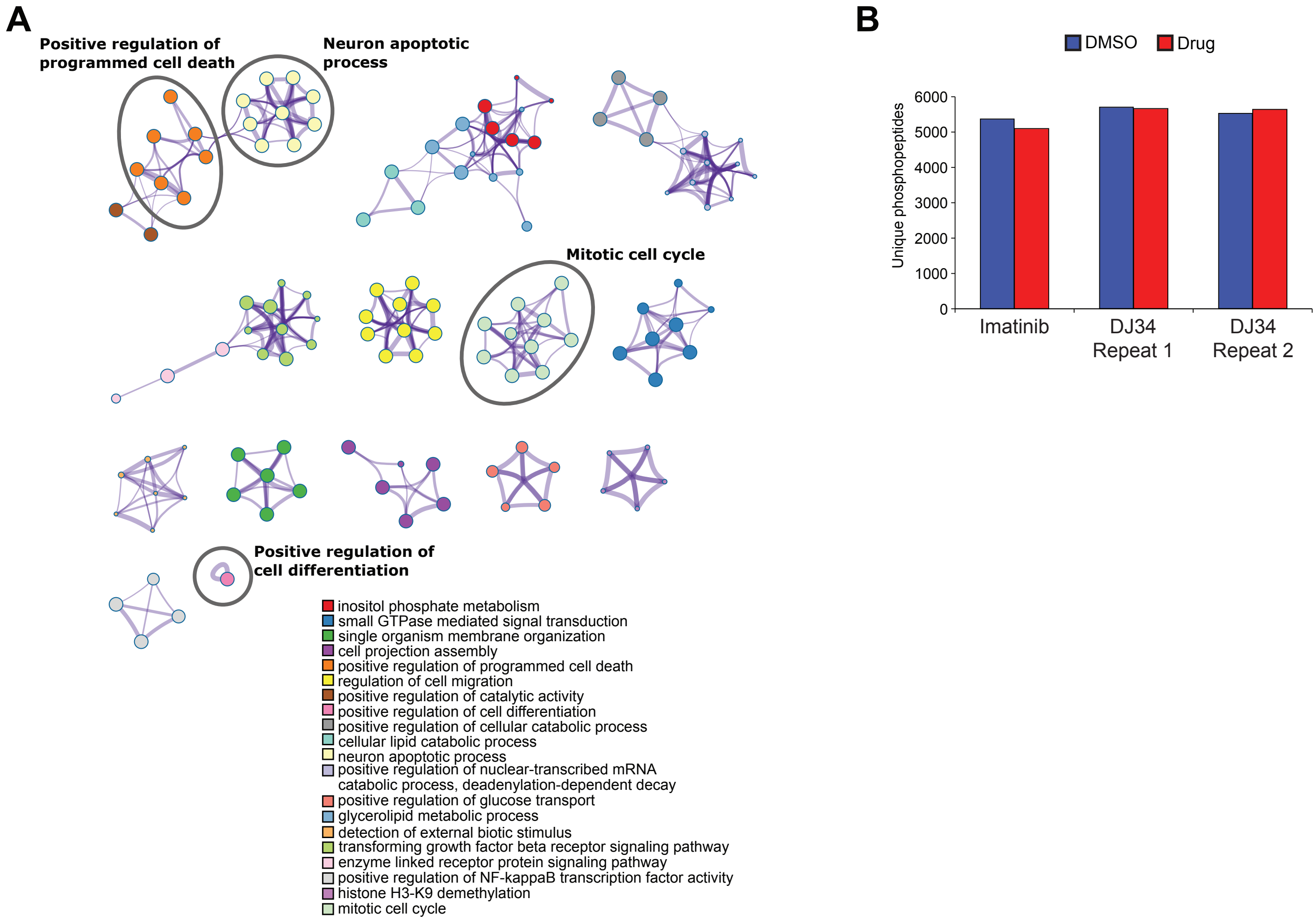

### Supplementary Figure S7

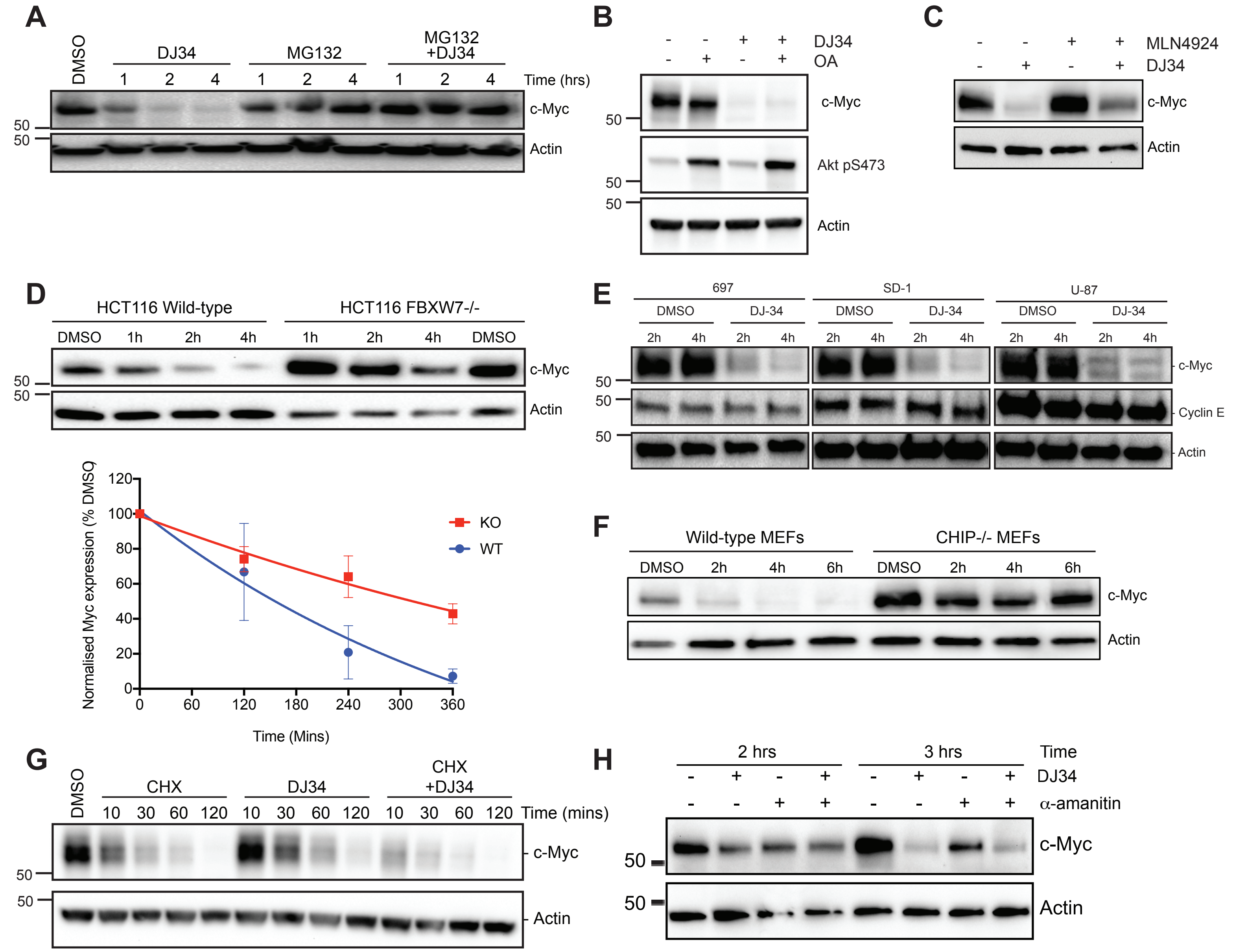


I

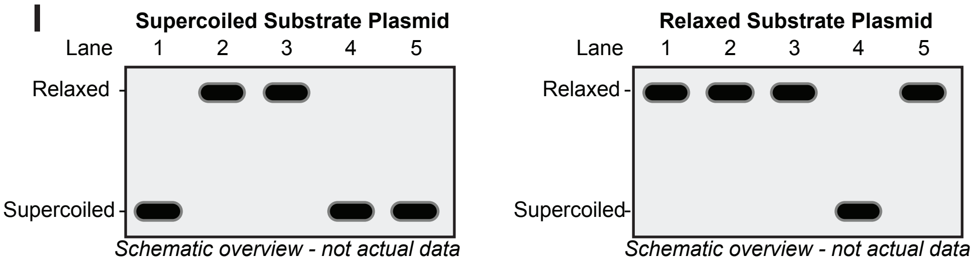


| | Present in the assay | | Effect of the compound | | |
|---|---|---|---|---|---|
| **Lane** | **Compound** | **Topo I** | **Intercalation** | **Topo I inhibition** | **Effect on supercoiled (S-plasmid) and Relaxed (R-plasmid) substrate** |
| 1 | DMSO (control) | - | N/A | N/A | Substrate-only negative control: No effect |
| 2 | DMSO (control) | + | N/A | N/A | Topo I positive control: S-plasmid becomes relaxed, R-plasmid remains relaxed |
| 3 | Compound A | + | - | - | No effect of the compound: S-plasmid becomes relaxed, R-plasmid remains relaxed |
| 4 | Compound B | + | + | - | S-plasmid remains supercoiled and R-plasmid also becomes supercoiled |
| 5 | Compound C | + | + | + | S-plasmid remains supercoiled and R-plasmid remains relaxed |

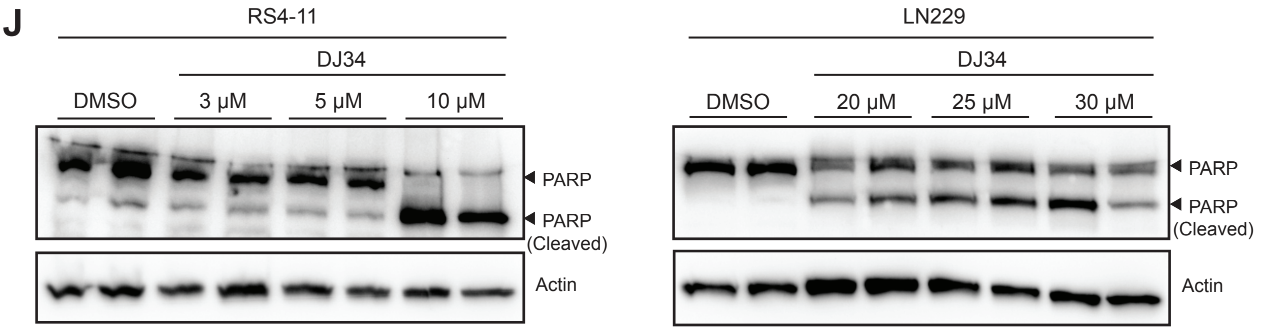

#### Supplemental Figure S8

Uncropped Westerns shown in Figure 4

Fig 4H: Myc pS62

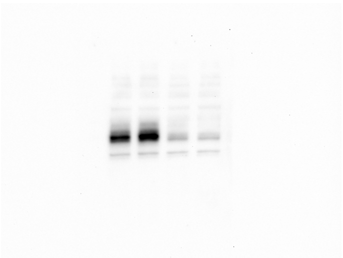

Fig 4H: Myc pT58

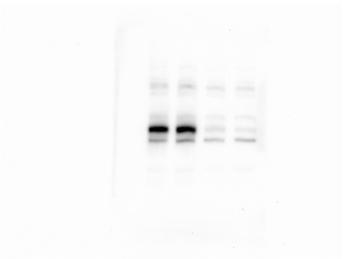

Fig 4H: Myc

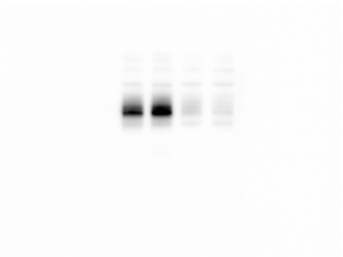

Fig 4H: p53 pS15

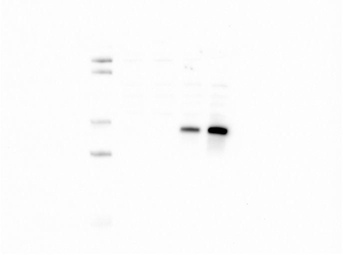

Fig 4H: p53

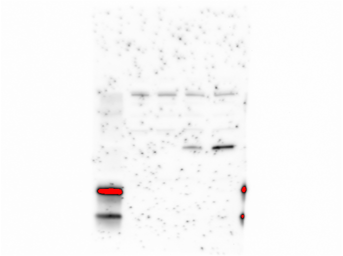

Fig 4H: Stat5



Fig 4H: Vinculin



Fig 4H: Actin

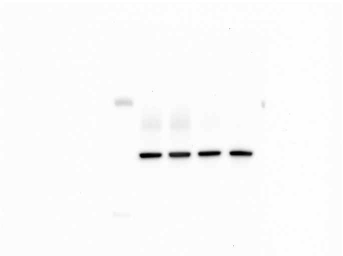

### Supplemental Figure S9

Uncropped Westerns shown in Figure 5

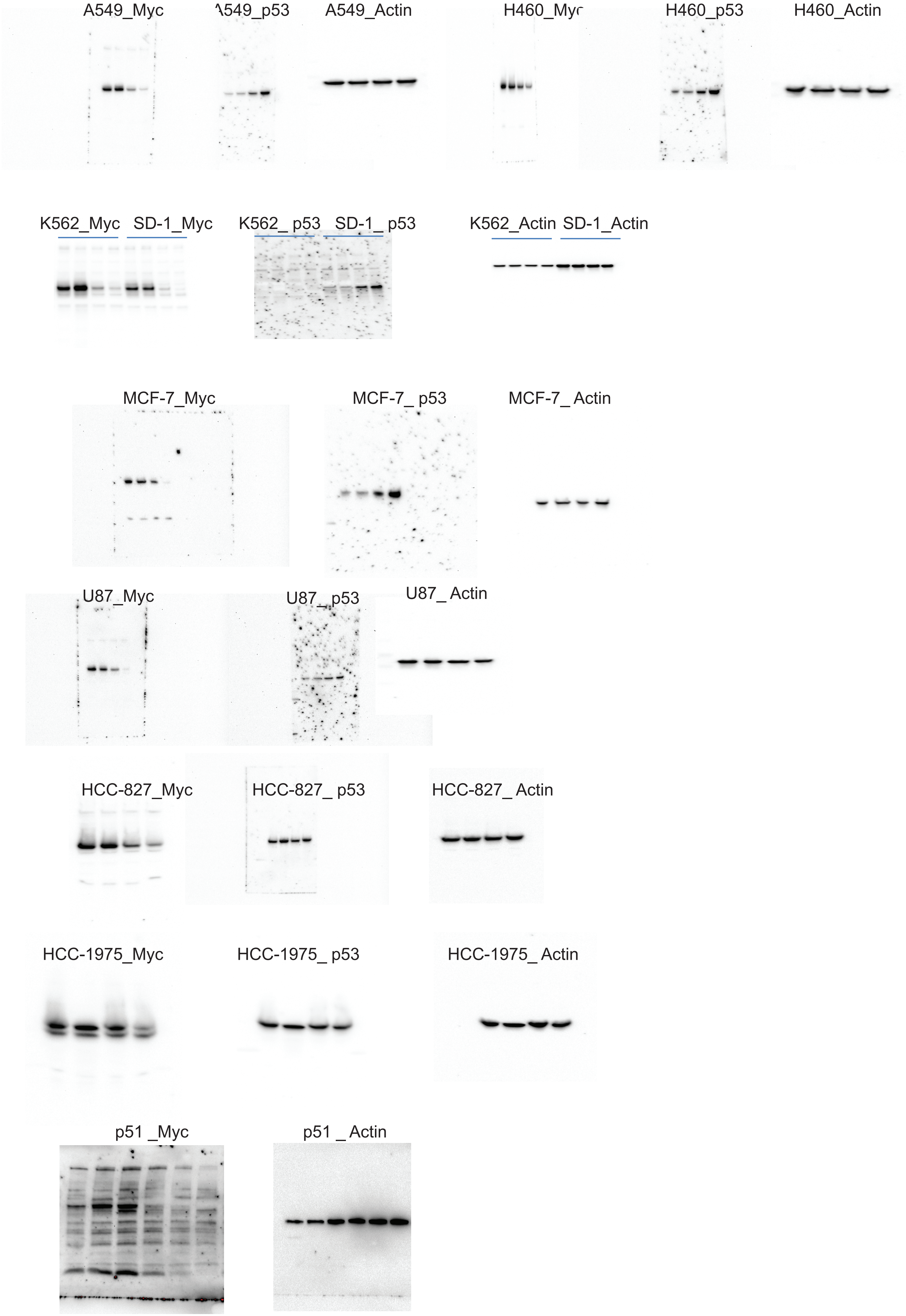

### Supplemental Figure S10

Uncropped Westerns shown in Figure 7

Fig 7B - K562 - c-Myc

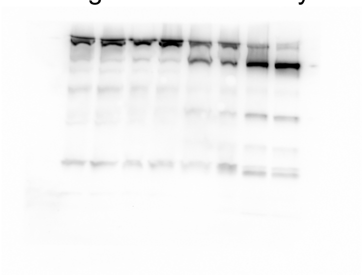

Fig 7B - MV4-11 - c-Myc

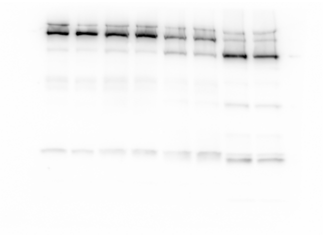

Fig 7B - K562 - Beta-Actin

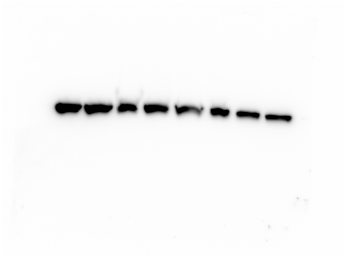

Fig 7B - MV4-11 - Beta-Actin

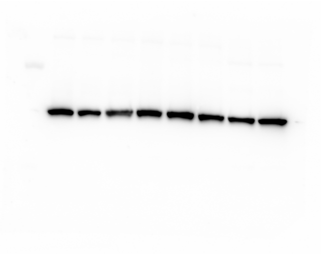

#### Supplemental Figure S11

Uncropped Westerns shown in Suppl. Fig. S5

Fig S5B: Akt pS473

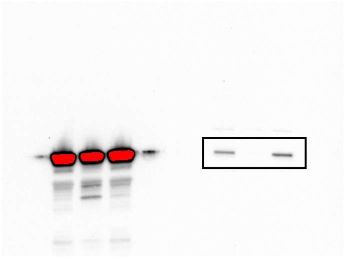

Fig S5B: Akt

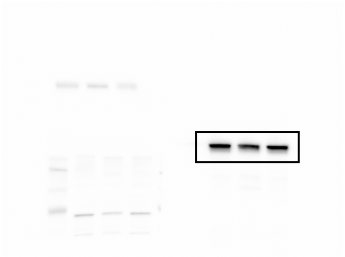

Fig S5B: Stat5 pY694

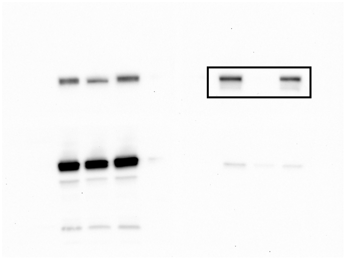

Fig S5B: Stat5

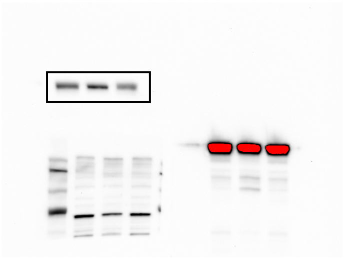

### Supplemental Figure S12

Uncropped Westerns shown in Suppl. Fig. S7

Fig S7B: Myc

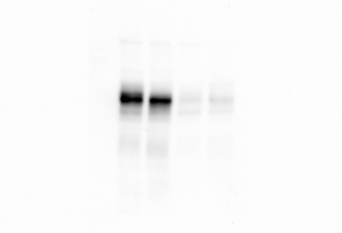

Fig S7B: Akt pS473

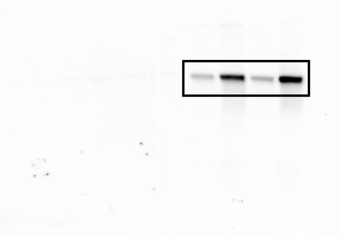

Fig S7B: Actin

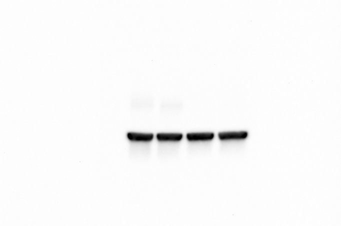

Fig S7C - c-Myc

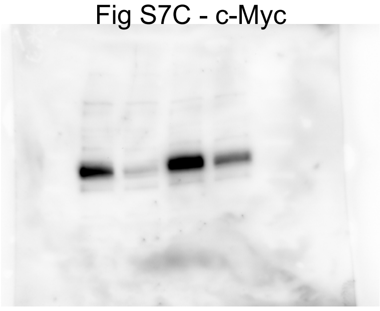

Fig S7C - Beta-Actin

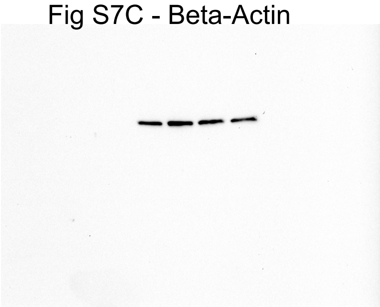

Fig S7D - c-Myc

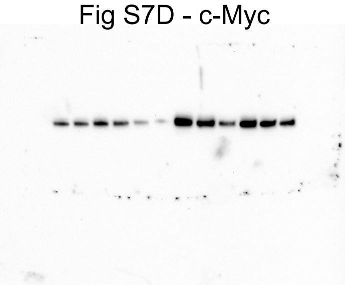

Fig S7D - Beta-Actin

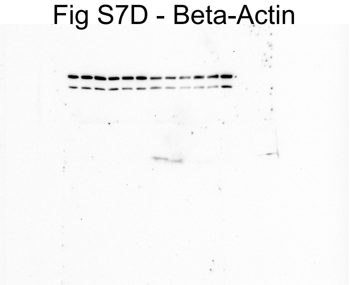

Fig S7F - c-Myc

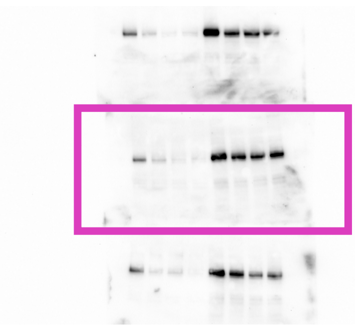

Fig S7F - Beta-Actin

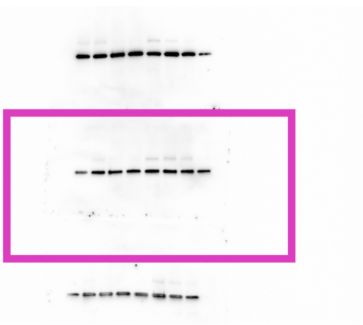

Fig S7G: Myc

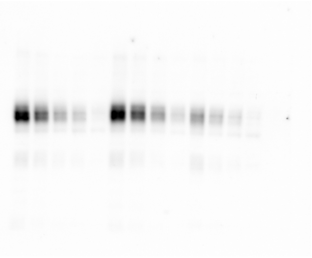

Fig S7G: Actin

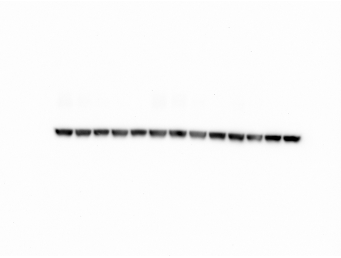

Fig S7H - c-Myc

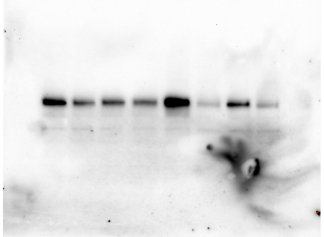

Fig S7H - Beta-Actin

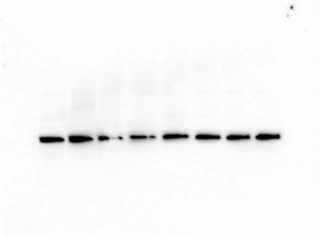

Fig S7J - RS4-11 - c-Myc

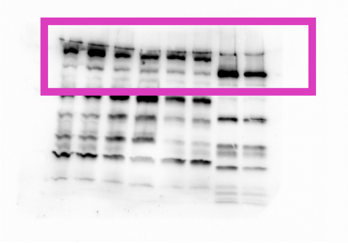

Fig S7J - LN229 - c-Myc

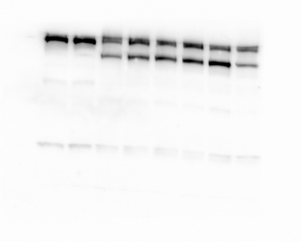

Fig S7j - RS4-11 - Beta-Actin

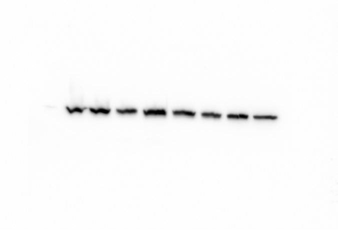

Fig S7J - LN229 - Beta-Actin

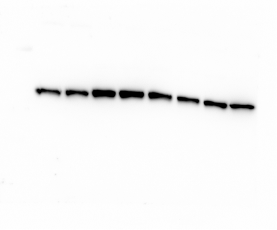
